# From urban runoff to mosquito success: spatiotemporal microbial assembly in larval water habitats under anthropogenic stressors

**DOI:** 10.64898/2026.06.01.729185

**Authors:** A. Gentil, E. Martin, L. Vallon, I. Tsirikhova, J. Gervaix, A. Fildier, L. Wiest, E. Vulliet, A. A. M. Cantarel, R. Cazabet, P. Luis, C. Valiente Moro

## Abstract

Urban mosquito habitats are heterogeneous aquatic ecosystems where anthropogenic inputs shape physicochemical conditions and microbial community assembly. However, the combined effects of environmental chemistry and microbial dynamics on mosquito fitness remain poorly understood across space and time. Here, we integrated environmental chemistry, metabarcoding, and experimental assays to investigate how spatiotemporal variation in urban larval habitats influences environmental microbial assembly and the biology of the Asian tiger mosquito, *Aedes albopictus*. Six stormwater drains were monitored over five months to characterize ions, dissolved gases, micropollutants, and bacterial and fungal communities. Laboratory assays using water from three contrasting habitats were then conducted to evaluate oviposition preference and mosquito performance. Microbial community composition was strongly structured by breeding-site identity and associated with distinct physicochemical signatures. Bacterial communities remained relatively stable over time, whereas fungal assemblages exhibited stronger temporal turnover. These environmental differences translated into marked variation in larval performance, ranging from rapid development and high survival to delayed development, reduced survival, and episodic cohort collapse under environmentally unstable conditions. Adult traits further revealed carry-over effects of larval environment exposure across life stages. Correlation analyses showed that mosquito fitness was associated with both abiotic variables and microbial taxa linked to larval survival, development, emergence, and adult longevity. In contrast, oviposition preference remained consistently high across habitats despite strong differences in offspring performance, indicating a decoupling between habitat attractiveness and suitability. Overall, our results demonstrate that anthropogenic stressors shape microbial assembly in urban larval habitats, with cascading consequences for mosquito fitness and population dynamics.

## Introduction

Microorganisms are ubiquitous across environments and play a fundamental role in ecosystem functioning and global biogeochemical cycles [1–3]. In aquatic environments, microbial communities regulate nutrient cycling [4], organic matter degradation [5], and the production of bioactive compounds [6]. They form complex assemblages whose composition and function are shaped by abiotic factors such as temperature, nutrients, and pollution, as well as by biotic interactions including competition, predation, and host activity [7–9]. Such dynamics are particularly evident in small standing water bodies, whose limited buffering capacity and strong physicochemical variability drive marked differences in microbial diversity across habitats [10, 11]. In urban ecosystems, habitat fragmentation and heterogeneous anthropogenic inputs create a mosaic of small aquatic habitats characterized by contrasting environmental conditions, leading to strong spatial turnover in microbial community composition [12, 13].

Microorganisms constitute the energetic foundation of aquatic food webs, supporting the growth of aquatic organisms either directly as a food resource or indirectly through trophic pathways in which primary consumers depend on microbial production [14, 15]. In addition, a subset of these microorganisms can colonize host surfaces or internal tissues, forming associations that contribute to the host microbiota [16, 17]. The composition of holobionts, defined as hosts and their associated microbiota including bacteria, fungi, and other microbial eukaryotes, is strongly influenced by the surrounding microbial environment [18, 19]. These environmental microorganisms can, in turn, shape host development, physiology, and fitness, thereby linking environmental conditions to organismal performance through microbially mediated processes [20–22]. Such microbe-host interactions are especially relevant in holometabolous insects, in which aquatic larval stages are primarily responsible for resource acquisition and somatic growth, whereas adults focus on reproduction and dispersal across distinct ecological niches [23–25]. Environmental conditions experienced during larval development can thus have lasting effects on adult phenotypic traits, including development time, survival, and reproductive success [26–28].

Urban mosquitoes provide an excellent model to study how spatiotemporal variation in environmental chemistry and microbial communities within larval aquatic habitats shapes ecosystem dynamics and influences life-history traits and oviposition behavior. Larvae develop in small, often transient aquatic habitats, such as stormwater drains or artificial containers, where environmental conditions can change rapidly, simultaneously affecting abiotic and biotic components of the habitat ([29, 12, 30, 31]. These habitats are further shaped by female oviposition decisions, which can influence population dynamics, as females tend to select breeding sites that maximize offspring performance [32– 34]. These choices are partly guided by multiple environmental cues, including chemical signals, microbial metabolites, and the presence of conspecifics or competitors, which collectively reflect habitat quality and productivity [35–37]. In addition, the physicochemical and microbial properties of larval habitats strongly influence larval development, adult fitness, oviposition behavior, and vectorial capacity [38–41]. Yet, most studies have focused on single factors or time points, overlooking the temporal dynamics that shape mosquito-microbiota-environment interactions in urban environments. Here, we combined environmental chemistry, metabarcoding, and controlled experimental assays to track temporal dynamics in water chemistry and microbial communities across the mosquito season. We then evaluated how these changes impact larval development, adult life-history traits, and oviposition behavior in the Asian tiger mosquito *Aedes albopictus*, one of the most common mosquito species inhabiting urban environments.

## Materials and Methods

### Study sites and sampling design

Six stormwater drains (B1-B6) were monitored at two-week intervals throughout the mosquito season (July-November 2024) in Villeurbanne, France (**Fig. 1; Table S1)**. Restricting the study to a single habitat type within one municipality limited structural and environmental heterogeneity unrelated to temporal dynamics. At the beginning of the study, all breeding sites were colonized by *Ae. albopictus* larvae, and their presence was confirmed by morphological identification. Each breeding site was sampled ten times (T1-T10) throughout the study period (**Fig. 1**), following the protocol described in the Supplementary Material. At each sampling time point, pH and temperature were measured *in situ* with indicator strips and a thermometer, and the presence of *Ae. albopictus* larvae was recorded. To assess the effects of breeding-site composition on mosquito oviposition behavior and life-history traits, additional assays were conducted at four time points: T1 (July), T4 (September), T6 (October), and T8 (November). These assays were restricted to three breeding sites (B1-B3), for which an additional 1 L of water was collected in a DURAN^®^ graduated glass bottle. At T6, axenic individuals used for life-history trait measurements became contaminated. Consequently, a new water sample was collected, creating an intermediate sampling point (T6.5) for breeding sites B1-B3, which was used exclusively for life-history trait measurements (**Table S1**).

**Figure 1.**
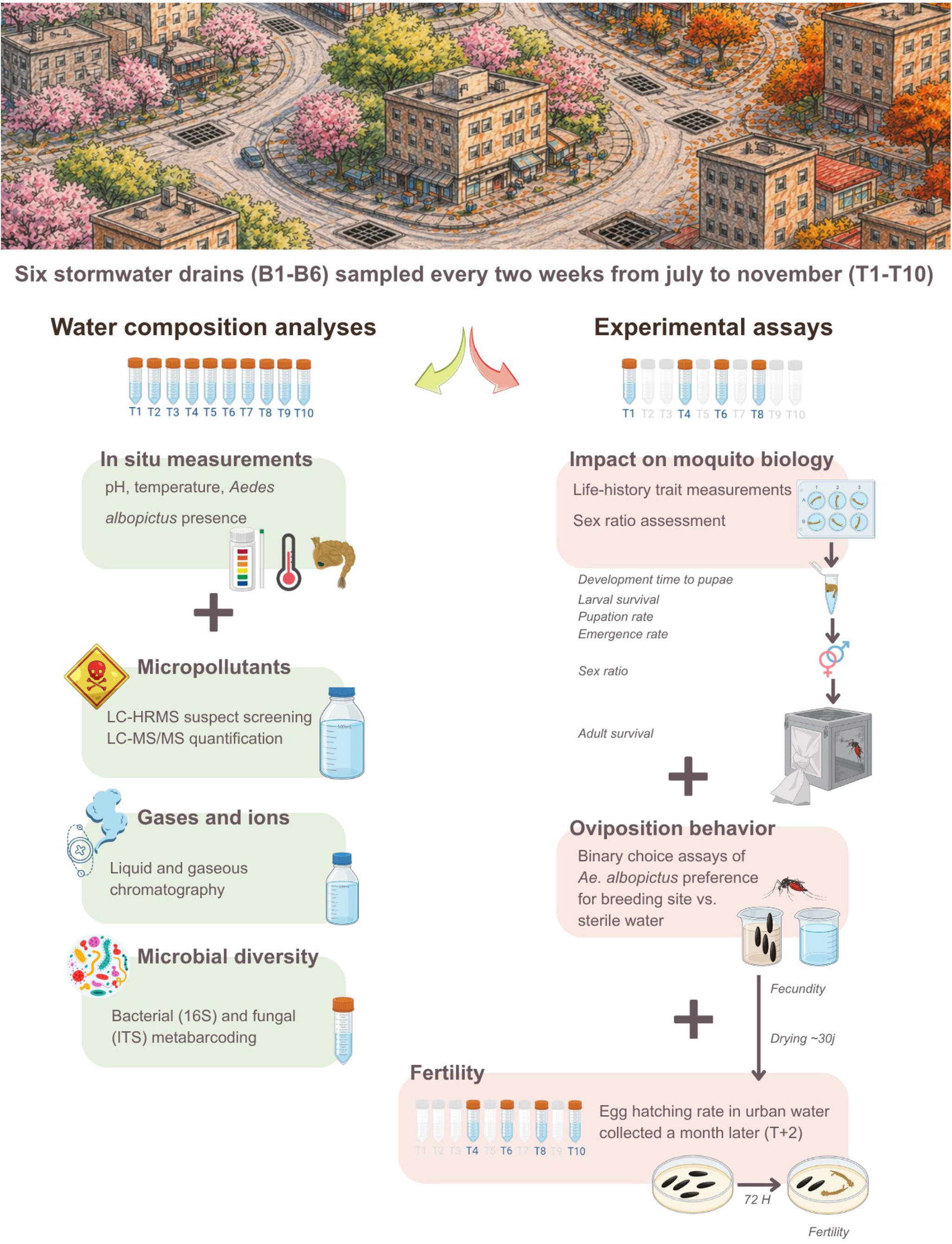
Overview of the experimental workflow. Water samples were collected every two weeks from July to November (T1-T10) at six urban breeding sites (B1-B6), all corresponding to stormwater drains colonized by Ae. albopictus larvae at T1. At each sampling time point, physicochemical parameters and microbial composition were characterized. Temperature and pH were measured in situ, along with the presence or absence of Ae. albopictus larvae. Analyses of ions and dissolved gases were performed ex situ using liquid and gas chromatography, respectively. Micropollutants and microbial diversity were also assessed ex situ throughout the study using high-resolution mass spectrometry (HRMS) and metabarcoding approaches targeting the 16S rRNA gene and fungal ITS regions, respectively. At sampling time points T1, T4, T6, and T8, additional water was collected from breeding sites B1, B2, and B3 to assess life-history traits and oviposition behavior in a laboratory-reared population of Ae. albopictus. To evaluate the impact of breeding site water composition on mosquito life-history traits while excluding microbial contamination not originating from the breeding sites, axenic larvae were obtained through successive washes of eggs with ethanol and bleach. After hatching, ten first-instar larvae were transferred under sterile conditions into each well of a six-well plate, with five plates per breeding site. This experiment was conducted in parallel with oviposition assays. Control treatments consisted of axenic larvae exposed either to sterile water supplemented with colony rearing tray water or to sterile water alone. Larvae were monitored daily to record survival and development time to pupation. Each pupa was then transferred to a 2 mL Eppendorf tube to allow for adult emergence and sex determination. Adults were monitored every three days for two months to assess potential carry-over effects of breeding site composition on adult survival. To evaluate the effect of breeding site composition on oviposition behavior and fertility, females aged 4-9 days were deprived of sugar for 24 h prior to artificial blood feeding on prewarmed sheep blood. Each gravid female was then individually placed in an experimental cage and offered a choice between two oviposition glass beakers: one containing water from an urban breeding site (B1, B2, or B3) and the other containing sterile water (control). Females (n = 24 per experiment) were allowed to oviposit for four days. Oviposition preference was assessed by counting the number of eggs laid in each beaker. Fecundity was estimated as the total number of eggs laid per female. Eggs were then dried at 28 °C in sterile Petri dishes for up to one month before being re-immersed in urban water collected at T+2 to assess hatching success (fertility).

### Physicochemical characterization of breeding site waters

Water samples were analyzed for dissolved gases and inorganic ions following the analytical protocols described in Duval et al. (2024), with detailed methodological descriptions provided in the Supplementary Material. For micropollutant analysis, water samples were filtered, spiked with isotopically labelled internal standards, and concentrated using a vacuum-assisted parallel evaporation system. Compounds were identified by UHPLC–HRMS using suspect screening with reverse-phase and HILIC separations. Targeted quantification was then performed by UHPLC–MS/MS in MRM mode. Full analytical procedures and instrumental settings are provided in the Supplementary Material.

### Environmental DNA extraction, metabarcoding, and sequence processing

Bacterial and fungal communities in breeding site waters were characterized by environmental DNA (eDNA) metabarcoding, with eDNA extracted from 8 mL of water per sample following Duval et al. (2024), with full methodological details provided in the Supplementary Material. Extraction blanks were included in each batch, and DNA concentrations were quantified using the Qubit dsDNA High Sensitivity assay (Thermo Fisher Scientific). Bacterial diversity was assessed by amplifying the V5-V6 region of the 16S rRNA gene using primers 784F/1061R (Andersson et al. 2008), and fungal diversity by targeting the ITS2 region using the gITS7/ITS4 primers [42]. Amplicon libraries were prepared using a two-step PCR approach to incorporate Illumina adapter sequences, as described by Duval et al. (2024). Negative controls (blank extractions) were included throughout. The second PCR step and paired-end sequencing (2 × 300 bp) were performed by Microsynth (Switzerland) on an Illumina MiSeq platform. High-throughput sequencing data were processed using the FROGS pipeline on the Galaxy platform following a workflow adapted from Antonelli et al. 2025 [39] and detailed in the Supplementary Material.

### Oviposition assays and life-history trait measurements

Behavioral assays and life-history trait analyses were conducted using the F27 and F28 generations of the *Ae. albopictus* laboratory colony (*AealbVB*), reared as described in the Supplementary Material. Oviposition preference assays were conducted using water collected from three urban breeding sites (B1-B3) at four sampling time points (T1, T4, T6, and T8) corresponding to monthly intervals between July and November (**Fig. 1**). Prior to experiments, water samples were filtered through filter paper to remove organic debris as well as conspecific larvae and eggs, which can influence oviposition through chemical cues. The experimental design is described in detail in the Supplementary Material. At the end of the exposure period, oviposition papers were collected and stored individually in sterile Petri dishes. Remaining water from each beaker was filtered to recover eggs laid directly on the water surface. Papers were dried and maintained in a climatic chamber for up to one month before egg counting. Females that died during the assay or did not lay eggs were excluded from subsequent analyses. In addition to oviposition assays, and to prevent interference from environmental microorganisms not naturally present in the tested waters, the effects of urban breeding site waters on mosquito life-history traits were assessed using axenic larvae following [41]. Full methodological details are provided in the Supplementary Material, along with descriptions of all life-history trait measurements.

### Statistical analyses

All statistical analyses were performed in R (v4.4.1). Abiotic and microbial variation among breeding sites and sampling time points was assessed using principal component analysis (PCA), non-metric multidimensional scaling (NMDS), diversity metrics, and permutational multivariate analysis of variance (PERMANOVA). Mosquito oviposition behavior and life-history traits were analyzed using generalized linear models, generalized linear mixed-effects models, beta-regression, and survival models, with breeding site, sampling time point, and their interaction included as explanatory variables. Associations between environmental parameters, microbial communities, and mosquito traits were explored using Spearman’s rank correlation analyses. Model selection was based on Akaike’s Information Criterion (AIC), and pairwise comparisons were adjusted using Tukey’s post hoc tests. Further details on statistical analyses are provided in the Supplementary Material.

## Results and discussion

### Environmental heterogeneity structures microbial assembly and physicochemical variation across urban mosquito breeding sites

We characterized biotic and abiotic properties of six urban mosquito breeding sites (B1-B6) across ten sampling time points spanning seasonal variation. Physicochemical analyses detected thirteen micropollutants including pharmaceuticals (paracetamol), biometabolites (adenine and nicotinamide), pesticides (carbendazim, diuron, monuron, fluopyram, terbutryn), repellents (DEET, icaridin), and nicotine-related molecules (nicotine, cotinine, hydroxycotinine), together with major ions, dissolved gases, pH, and temperature (**Fig. 2**). Despite their shared urban context, breeding sites exhibited strongly contrasted physicochemical profiles (**Fig. 2A**), indicating that each breeding site constituted a distinct environmental niche. B3 showed the most chemically enriched signature, characterized by elevated ion and micropollutant concentrations as well as higher temperature and more alkaline conditions (**Fig. 2A**). In contrast, B1 exhibited lower micropollutant contamination but elevated dissolved gas concentrations. B4 displayed recurrent peaks of repellents and pharmaceuticals, likely reflecting localized human activity. B5 and B6 presented overall lower chemical concentrations (**Fig. 2A**). Heatmap visualization of chemical variables revealed clear differences across all breeding sites in both the occurrence and concentrations of micropollutants, as well as in ionic composition and dissolved gases (**Fig. 2 B-D**). Several compounds displayed site-specific or episodic peaks across sampling dates, suggesting episodic anthropic inputs associated with urban activities including traces of domestic wastewater, runoff from surrounding surfaces, or direct deposition associated with the urban use of repellents, pharmaceuticals, and pesticides [43]. Consistently, the abiotic PCA showed that the first two axes explained 29.0% and 12.1% of the variance, respectively, with samples clustering primarily by breeding site, along with some temporal separation and only limited segregation according to colonization status (**Fig. 3, Table 2**). One-factor PERMANOVAs supported these patterns, showing significant effects of breeding site identity, sampling time point, and colonization status on abiotic profiles (*p* = 0.001). While some studies on *Ae. albopictus* have reported significant associations between larval occurrence and specific variables such as pH or salinity, others highlight the species’ ability to colonize a wide range of water conditions, suggesting that abiotic factors alone may not reliably predict breeding-site colonization [44, 45]. Metabarcoding revealed highly diverse microbial assemblages within these human-made aquatic habitats, with 26,042 bacterial (16S rRNA gene) and 15,883 fungal (ITS) OTUs. Although the total number of OTUs recovered was high, richness per sample remained within the range reported for *Aedes* spp. urban water habitats using Illumina metabarcoding (e.g. ∼10^2^-10^3^ taxa per sample) (**Fig. 4**) [30, 46, 47]. Microbial community composition was primarily structured by breeding-site identity, although bacteria and fungi exhibited contrasting temporal dynamics (**Fig. 3; Table 2**). For bacterial communities, NMDS ordination (stress = 0.232) and PERMANOVA revealed a strong site effect on beta diversity (R^2^ = 0.375, *p* = 0.001), whereas temporal variation was not significant and colonization status had only a weak effect (R^2^ = 0.027, *p* = 0.020) (**Fig. 3, Table 2**). Consistently, bacterial richness and Shannon diversity varied significantly among breeding sites (*p* < 0.001) but remained stable across sampling time points, with only a marginal effect of colonization status on richness (*p* = 0.0499), and no effect on Shannon or Simpson diversity (*p* = 0.148 and *p* = 0.260, respectively) (**Fig. 4, Table 2**). Together, these results indicate that bacterial communities remained stable once established, potentially reflecting functional redundancy and rapid turnover buffering short-term environmental fluctuations [48]. Fungal communities displayed somewhat different dynamics. NMDS ordination (stress = 0.202) and PERMANOVA likewise showed a strong effect of breeding-site identity on fungal beta diversity (R^2^ = 0.354, *p* = 0.001), together with a weaker temporal signal (R^2^ = 0.183, *p* = 0.070) and a significant effect of colonization status (R^2^ = 0.031, *p* = 0.014) (**Fig. 3, Table 2**). In contrast to bacteria, fungal richness and Shannon diversity varied significantly across sampling time points (*p* < 0.01), in addition to strong site effects (*p* < 0.001) (**Fig. 4, Table 2**). Colonization status also influenced fungal richness and Shannon diversity (*p* < 0.01). Together, these results indicate that fungal communities were more temporally dynamic and environmentally responsive than bacterial communities, consistent with a previous study showing greater fungal sensitivity to environmental heterogeneity in mosquito larval habitats [49]. More broadly, fungal communities in aquatic ecosystems are often more responsive to environmental heterogeneity, likely reflecting differences in dispersal, life-history traits, and sensitivity to local conditions [50]. Larval colonization also emerged as a key driver of microbial composition, likely due to multiple interactions between larvae and their aquatic habitat. By ingesting microorganisms and organic matter, larvae exert top-down grazing pressure that alters microbial abundance and selects for specific taxa, while larval excretions modify nutrient availability and water chemistry, further shaping microbial assemblages [51, 52]. This site structuring was also evident in taxonomic composition. Although metabarcoding limits fine-scale taxonomic resolution, family-level analyses revealed strong site specificity rather than uniform responses to contamination gradients (**Fig. 5**). In particular, B3, characterized by the most contrasting physicochemical profile in terms of ions and micropollutants (**Fig. 2**), harbored the lowest bacterial richness (**Fig. 4**) and a distinct microbial assemblage (**Fig 5**) with several taxa reduced or absent relative to other breeding sites **(Fig 5**). Moreover, breeding sites with comparable alpha diversity (e.g., B1 and B2) differed substantially in taxonomic composition, highlighting that similar richness does not necessarily imply similar community structure [53, 54]. To further explore potential links between environmental conditions and microbial communities, we examined correlations between microbial families and physicochemical variables. Few strong one-to-one associations were detected (**Fig. 6**). Instead, most microbial families showed moderate correlations with multiple ions and micropollutants, suggesting that microbial assemblages were shaped by multivariate environmental filtering rather than by individual compounds alone. This pattern is consistent with the chemical complexity of urban waters exposed to mixtures of anthropic contamination [55], as well as with experimental evidence showing that pollutant mixtures can exert stronger ecological effects than single compounds [56, 57]. Together, these results highlight the importance of considering combined environmental stressors when interpreting microbial community assembly in anthropogenic habitats. Because mosquito larvae acquire most of their microbiota horizontally from the aquatic environment through feeding and filter activity, the environmental microbial pool is a key determinant of host-associated community assembly [58, 59]. Thus, chemical mixtures may influence mosquito-associated microbiota indirectly by restructuring environmental microbial communities rather than selectively eliminating individual taxa [60]. Given that larval development depends on microorganisms both as a food resource and as developmental cues [41, 61], persistent differences in microbial richness and composition across sites may affect larval performance, development time, vector competence, and breeding-site attractiveness [62, 40]. In this context, chemically enriched habitats such as B3, characterized by high pesticide loads and reduced microbial richness, illustrate how chronic pesticide exposure may alter microbial community structure and functional potential [63, 64], potentially modifying both the microbial taxa available for larval colonization and the resources supporting larval growth. For subsequent analyses, we focused on three pre-selected sites (B1, B2, B3) monitored over time, with environmental conditions ranging from permissive to chronically constrained, and chemically unstable conditions, allowing us to investigate their potential effects on mosquito life-history traits and oviposition preferences.

**Figure 2.**
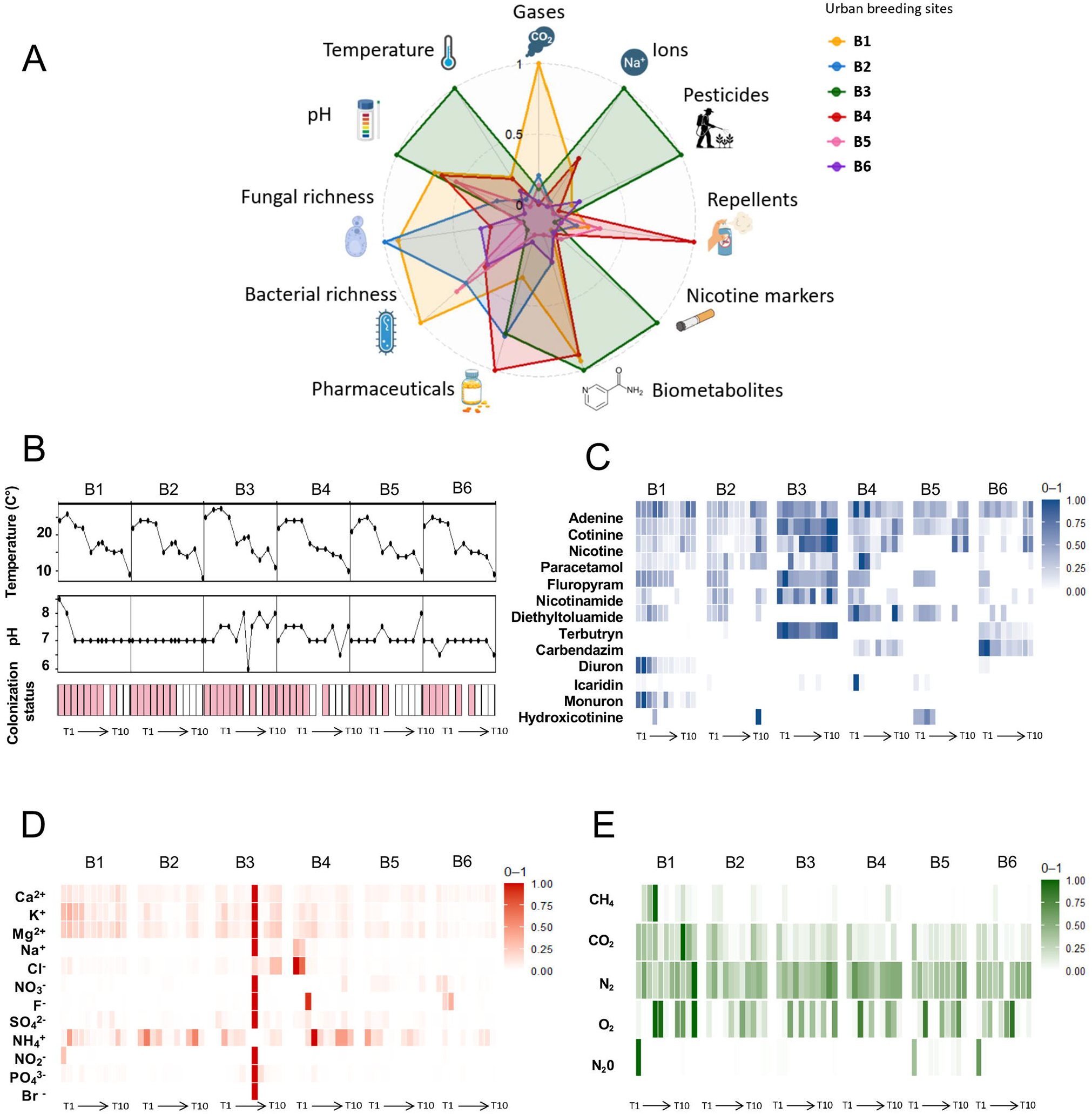
Temporal variation in physicochemical properties, microbial composition, and colonization status across urban mosquito breeding sites. (A) Mean normalized values (0-1) across all sampling time points (T1-T10) for each breeding site (B1-B6) describing both biotic (fungal and bacterial richness) and abiotic properties (pH, temperature, pesticides, repellents, nicotine markers, biometabolites, pharmaceuticals, dissolved gases and ions). (B) Water temperature, pH, and colonization status of each breeding site (B1-B6) over time (T1-T10), with pink color indicating the presence of larvae. (C) Micropollutants detected and quantified across breeding sites (B1-B6) over time (T1-T10), with concentrations expressed in µg/L and shown as normalized values (0-1). (D) Ions detected and quantified across breeding sites (B1-B6) over time (T1-T10), with concentrations expressed in µg/L and shown as normalized values (0-1). (E) Dissolved gases detected and quantified across breeding sites (B1-B6) over time (T1-T10), with concentrations expressed in µg/L and shown as normalized values (0-1).

**Figure 3.**
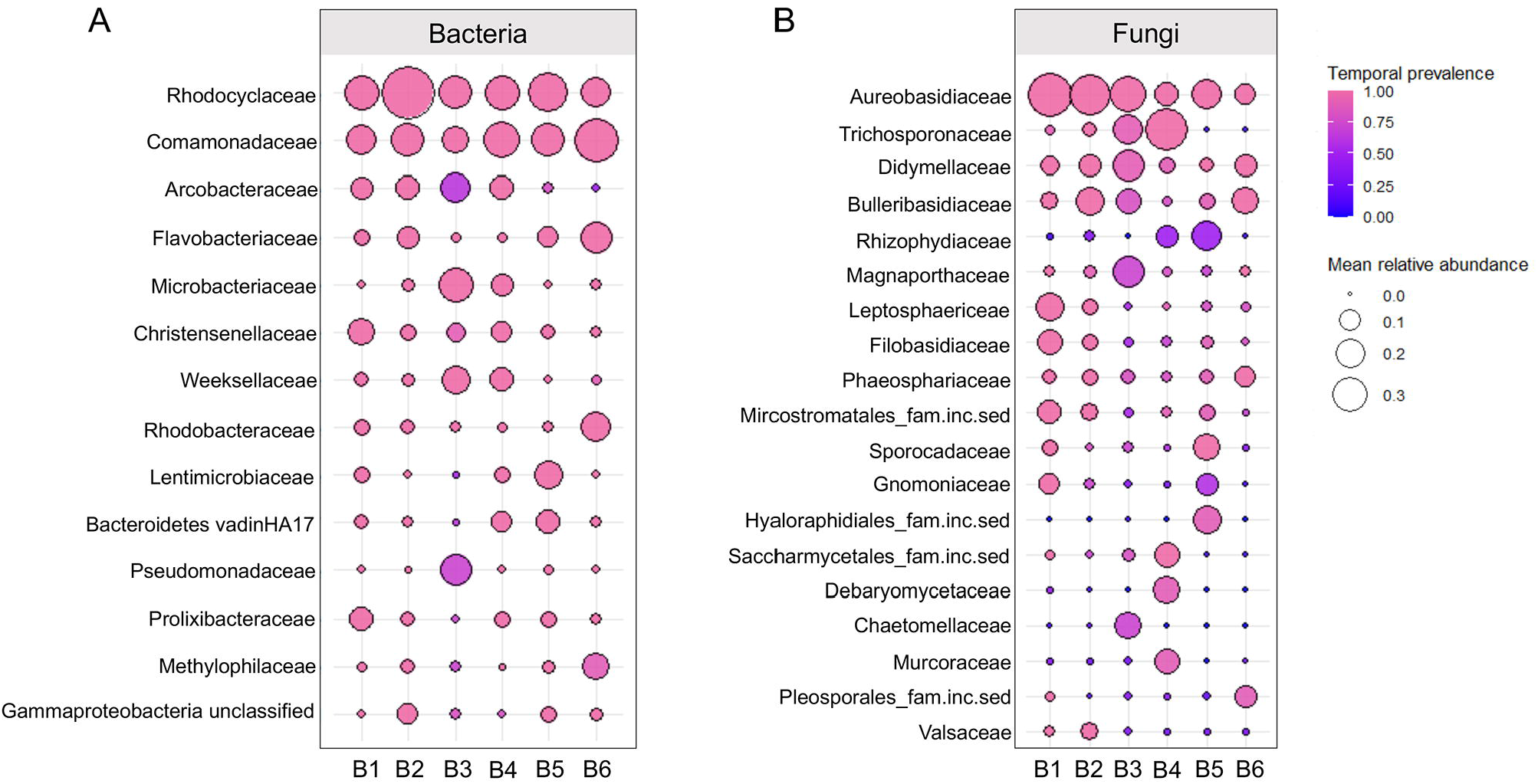
Representation of the most abundant microbial families across breeding sites, as identified through metabarcoding. (A) Bubble plot showing the most abundant bacterial families in each breeding site (B1-B6). (B) Bubble plot showing the most abundant fungal families in each breeding site (B1-B6). Bubble size represents mean relative abundance across the sampling period while color indicates temporal prevalence across the sampling period within each breeding site.

**Figure 4.**
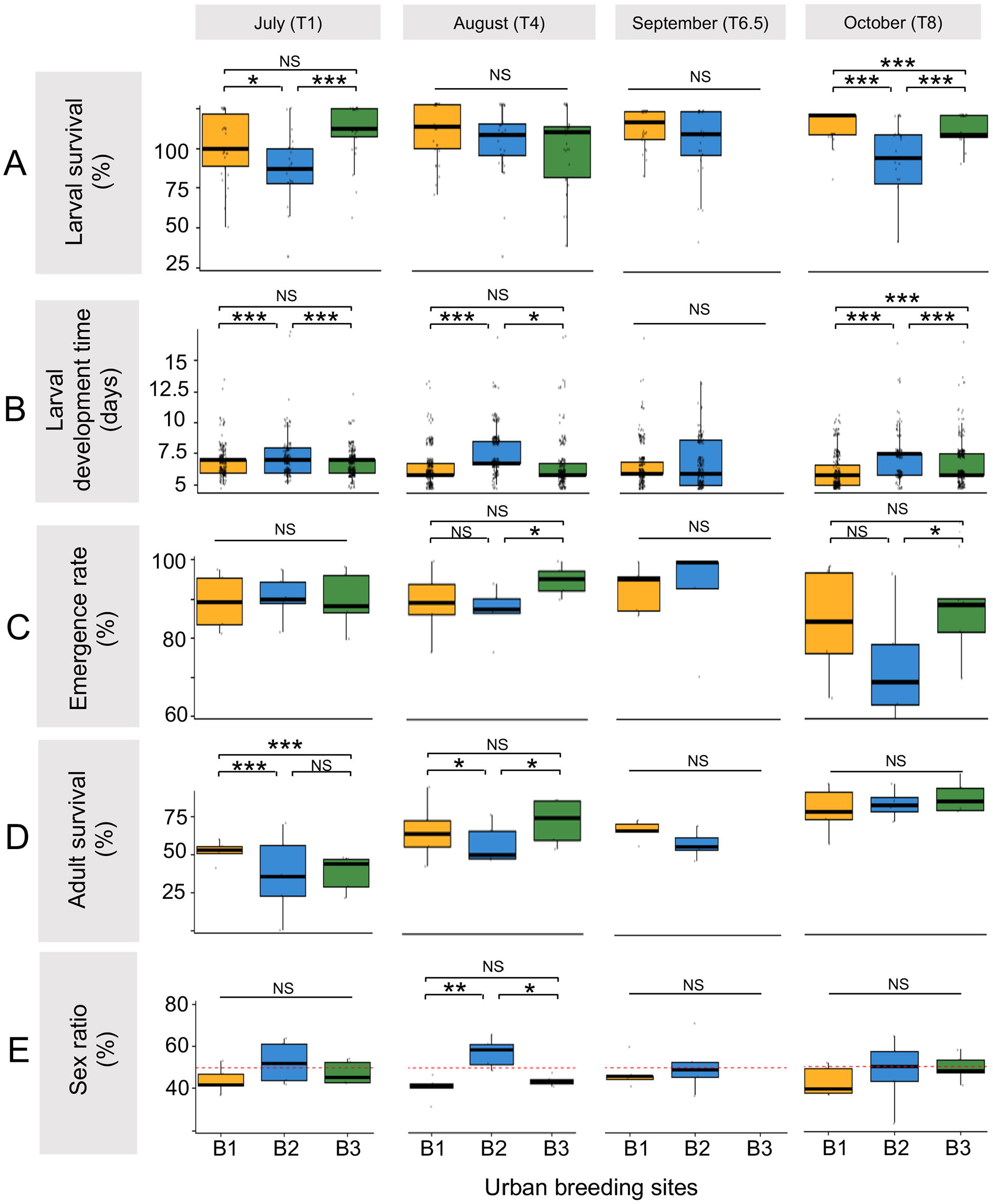
Temporal effects of urban breeding sites (B1-B3) on life-history traits of *Ae. albopictus* from larval development to adult survival. Each panel represents a specific life-history trait and is divided by experimental times (T1: July, T4: August, T6.5: September, T8: October) and breeding sites (B1, B2, B3). (A) Boxplots representing larval survival rate, in percentage (%), with the median indicated by a black line. (B) Boxplots representing development time to the pupal stage in days, with the median indicated by a black line. (C) Boxplots representing pupation rate in percentage (%), with the median indicated by a black line. (D) Boxplots showing adult survival at 35 days, in percentage (%). Individuals were monitored for up to 60 days, and the median is indicated by a black line. (E) Boxplots representing sex ratio (percentage of females) (%) in each cage, with the median indicated by a black line and the red dashed line indicating parity (50%). Larval development time was analyzed using accelerated failure time (AFT) models with right-censoring. Weibull, log-normal and log-logistic distributions were compared using the Akaike Information Criterion (AIC). The best-supported model included breeding site, sampling time point and their interaction, with pairwise comparisons estimated from marginal means (Tukey-adjusted). Larval survival, pupation probability and adult emergence were analyzed using binomial generalized linear mixed models (GLMMs) with breeding site, sampling time point, and their interaction as fixed effects, and well, plate, or cage as random effects when appropriate. Sex ratio was analyzed using binomial models of female and male counts per cage. As the cage random effect was negligible, final inference relied on a binomial GLM with Tukey-adjusted comparisons. Adult survival was analyzed using AFT models selected by AIC and confirmed with a Cox proportional hazards model accounting for cage clustering. For clarity, only differences between breeding sites within each sampling time point are shown in the figure. Temporal differences within each breeding site were also examined, and are reported in Table S9. Statistical significance is shown above the boxplots with “*” indicating p < 0.05, “**” indicating p < 0.01, “***” indicating p < 0.001, and “NS” indicating *p >* 0.05.

**Figure 5.**
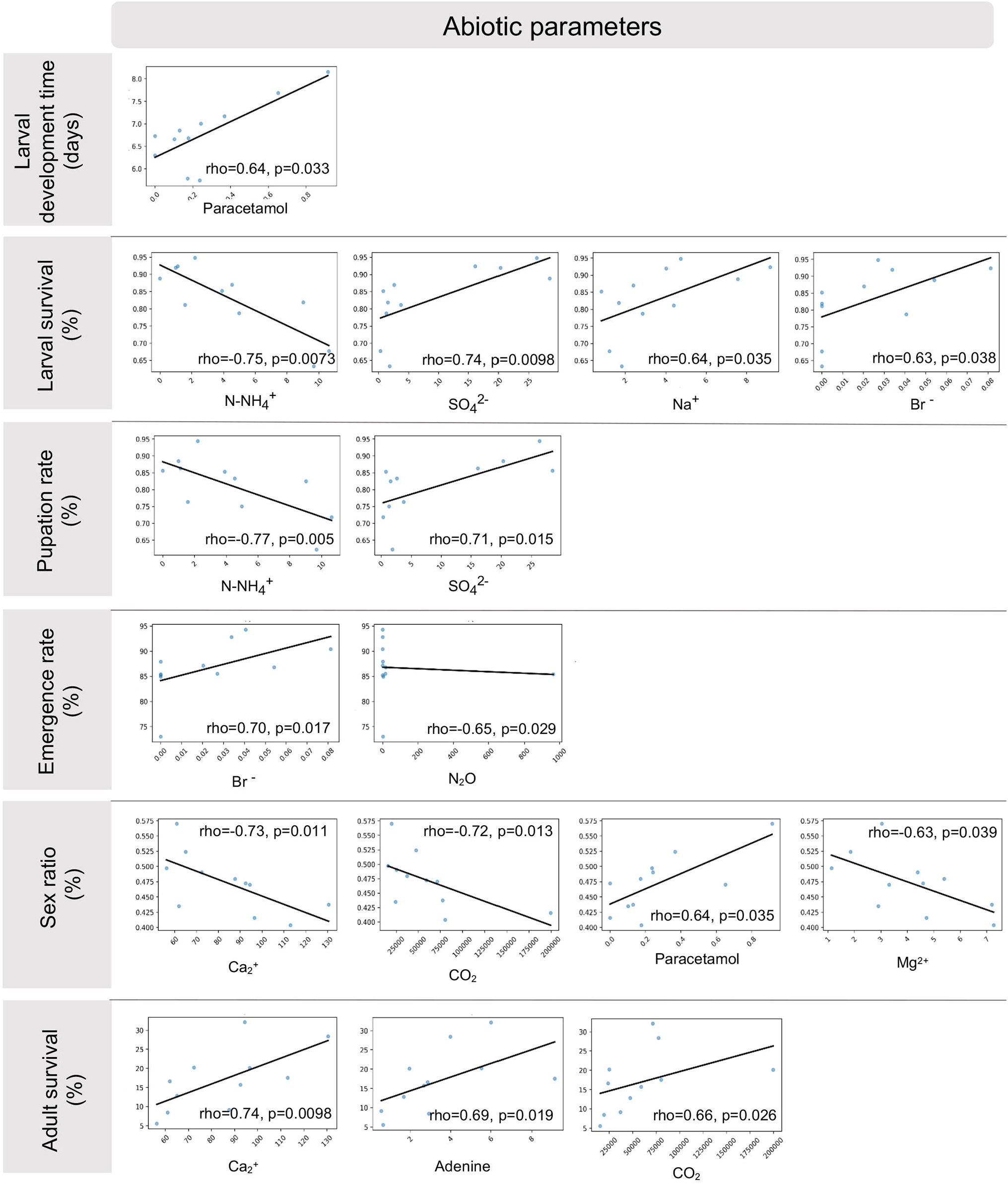
Significant correlations between abiotic parameters and mosquito life-history traits across urban breeding sites. Spearman correlation analyses were performed between measured abiotic parameters and Ae. albopictus life-history traits across experimental breeding sites (B1-B3) and sampling time points (T1, T4, T6.5, T8). Only statistically significant correlations are represented and a linear fit is plotted for readability. (A) Correlations involving larval survival rate (%). (B) Correlations involving larval development time to pupation (days). (C) Correlations involving pupation rate (%). (D) Correlations involving adult emergence rate (%). (E) Correlations involving adult survival at 35 days (%). (F) Correlations involving sex ratio (% females). Spearman’s correlation coefficients (rho) and associated p-values are indicated for each statistically significant association. Positive and negative relationships are represented by ascending and descending regression lines, respectively. Abiotic parameters included ions (NH_4_^+^, SO_4_^2−^, Na^+^, Br^−^, Ca^2+^, Mg^2+^), dissolved gases (CO_2_, N_2_O), and micropollutants (paracetamol and adenine). Only correlations with p < 0.05 are displayed.

**Figure 6.**
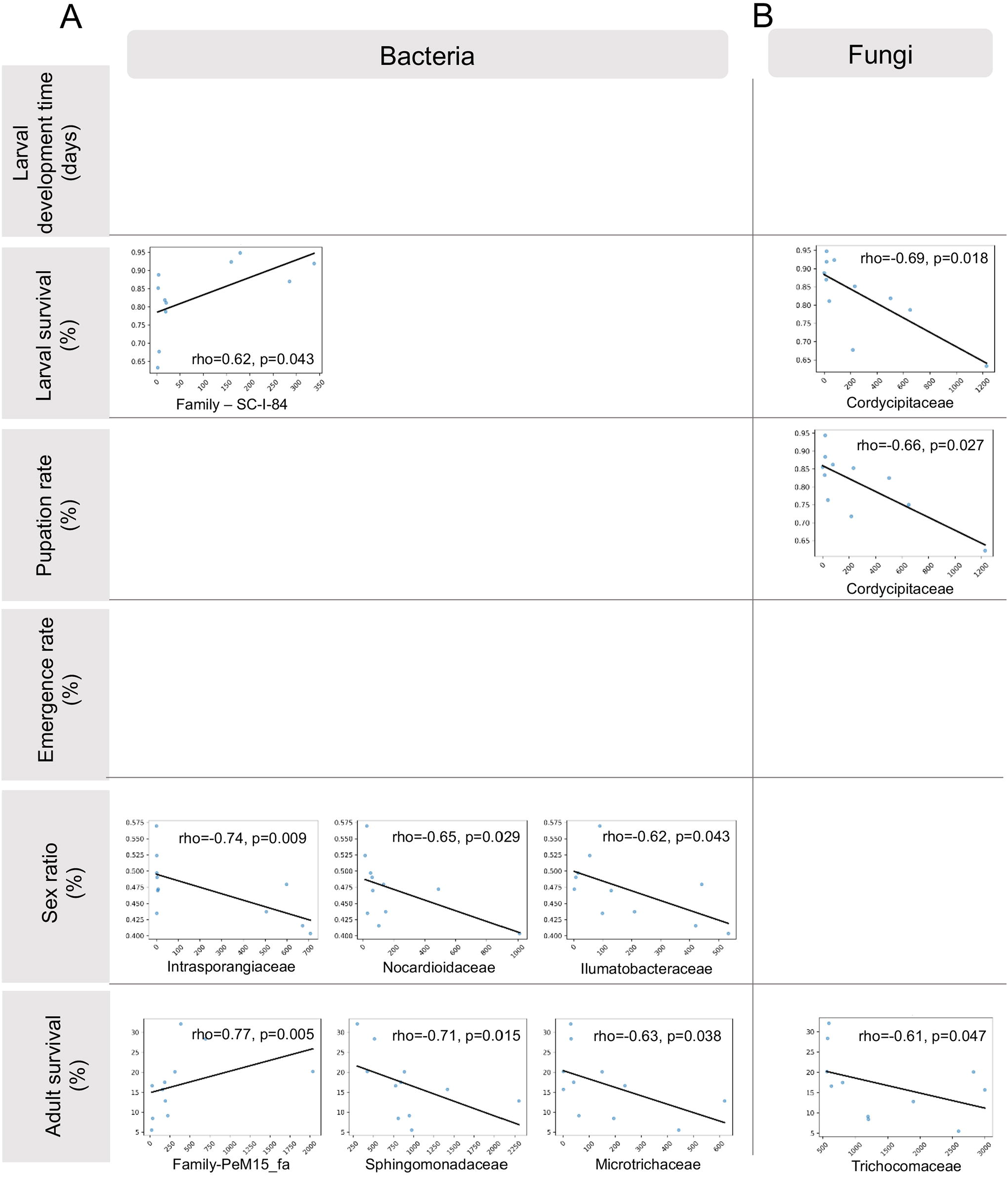
Significant correlations between dominant microbial families and mosquito life-history traits across urban breeding sites. Spearman correlations analyses were performed between the relative abundance of dominant bacterial and fungal families identified by metabarcoding and Ae. albopictus life-history traits across experimental breeding sites (B1-B3) and sampling time points (T1, T4, T6.5, T8). Only statistically significant correlations are represented. (A) Correlations involving bacterial families and mosquito life-history traits. (B) Correlations involving fungal families and mosquito life-history traits. Life-history traits include larval survival rate (%), larval development time to pupation (days), pupation rate (%), adult emergence rate (%), adult survival at 35 days (%), and sex ratio (% females). Spearman’s correlation coefficients (rho) and associated p-values are indicated for each statistically significant association. Positive and negative relationships are represented by ascending or descending regression lines, respectively. Bacterial families included Sphingomonadaceae, Microtrichaceae, Intrasporangiaceae, Nocardioidaceae, Ilumatobacteraceae, Family-SC-I-84, and Family-PeM15_fa, whereas fungal correlations mainly involved Cordycipitaceae and Trichocomaceae. Only correlations with p < 0.05 are displayed.

### Environmental heterogeneity drives mosquito performance but decouples oviposition from fitness outcomes

Larval development time differed significantly among breeding sites and over sampling time points (log-logistic AFT: χ^2^ = 191.97, df = 11, p < 0.001; **Fig. 7, Table 3**), with a significant site × month interaction indicating temporally dynamic environmental effects. Over the study period, B1 consistently supported the fastest development, with mean development time decreasing from 7–8 days in summer to 6.3 days in October (**Fig. 7**). In contrast, B2 exhibited persistently longer development times, with differences already detectable in summer (∼6.8–7.0 days vs slightly lower values in B1), and becoming more pronounced in October, when larvae required on average 7.7 days to complete development compared with 6.3 days in B1 and 7.0 days in B3 (B1–B2 estimate = −0.436, *p* < 0.0001). B3 displayed high performance under permissive conditions but experienced near-complete cohort collapse in September, preventing parameter estimation and highlighting acute stochastic instability. Survival analyses mirrored these patterns, with a strong site × month interaction (GLMM, binomial; **Fig. 7, Table 3**). B1 larvae showed robust and gradually improving survival, increasing from 79.9% in July to >90% in August and September, and reaching 95.1% in October. B2 larvae experienced consistently lower survival throughout the season, with reduced survival in July compared with B1 (OR = 2.35, *p* = 0.018), remaining lower in August, and reaching only 71.0% in October compared with 95.1% in B1 (OR = 9.57, *p* < 0.0001). B3 showed high survival during part of the season (comparable to B1 in July, August, and October; e.g., 92.3% in October), but experienced near-total mortality in September (0.4% survival), generating extreme statistical contrasts (*p* < 0.0001). Pupation probability was significantly influenced by site, month, and their interaction (GLMM, LRT p < 0.001; **Fig. 7, Table 3**). B1 pupation increased steadily over sampling time points, from 77% in July to approximately 88-92% in August–September, reaching 94.4% in October. B2 consistently exhibited lower pupation rates than B1, with significant differences in July (*p* = 0.0027) that persisted through August and September, and declined markedly in October (70.3% vs 94.4% in B1; *p* < 0.0001). This late-season drop paralleled the delayed development observed at this site. B3 showed strong temporal variability in pupation rates, with high values in July, a reduction in August relative to B1 (*p* = 0.0318), biologically uninterpretable values in September due to larval collapse, and a return to high levels in October (86.4%). Together, these trajectories indicate three distinct ecological regimes for larval development: (i) a permissive and progressively improving habitat (B1), (ii) a chronically constrained environment with cumulative late-season effects (B2), and (iii) a temporally unstable habitat alternating between permissive and disturbed states (B3). Importantly, B1 supported high larval performance despite the detection of site-specific herbicides, notably diuron and monuron. Although herbicides primarily target plants, low environmental concentrations can affect aquatic invertebrates, including mosquito species, through sublethal physiological stress, activation of detoxification pathways (e.g., cytochrome P450 enzymes), or indirect effects via microbial community shifts [39, 65–67]. By contrast, the reduced larval performance observed in B2 (slower development, reduced survival, and lower pupation) is more consistent with chronic environmental constraints than with acute toxicity. Unlike B1, which supported high performance despite detectable herbicides such as diuron and monuron, B2 did not display particularly elevated concentrations of micropollutants, ions, or dissolved gases. The main difference at this site was its higher fungal richness, suggesting that differences in microbial community structure, rather than chemical stress intensity alone, may contribute to reduced larval performance [68]. B3, in contrast, illustrates the importance of acute disturbance. The September collapse coincided with a high concentration of ions associated with low bacterial and fungal richness, suggesting a possible link with environmental disturbance, although the exact causes of larval mortality remain unresolved. Mosquito larvae are known to be sensitive to environmental contamination, as pollutants can impair survival and development through physiological or metabolic stress [69]. However, given the lack of direct evidence, B3 may be better described as a site subject to episodic or stochastic perturbations, rather than as consistently poor habitat quality. Overall, these results show that mosquito larval performance is structured by strong spatial and temporal environmental heterogeneity, reflecting interacting abiotic and biotic drivers rather than chemical conditions alone.

Adult performance metrics revealed site-specific carry-over effects from larval environmental exposure. Emergence success mirrored larval survival patterns, with slightly amplified contrasts among breeding sites (**Fig. 7, Table 3**). In B1, emergence remained consistently high throughout the study period (85.5-94.4%), indicating stable completion of development. In B2, emergence was moderate in July and August, comparable in September when larvae survived, but declined markedly in October (73%), suggesting cumulative late-season impairment. In B3, emergence was high when larval development was successful (94.3% in August and 90.4% in October), but no emergence occurred in September due to prior larval mortality. Adult survival revealed significant breeding-site-specific temporal dynamics (Weibull AFT: χ^2^ = 85.41, df = 11, *p* < 0.001; Cox regression interaction, robust *p* = 0.02; **Fig. 7, Table 3**). In July, survival at day 35 was highest in B1 (28.4%), significantly exceeding B2 (12.8%) and B3 (15.7%) (*p* < 0.001). B1 remained relatively stable in August (20%), while B2 and B3 increased slightly but did not surpass it. Notably, B2 displayed a marked increase in adult survival in October (32.1%) despite reduced larval performance, while B3 exhibited a collapse in September followed by recovery in October. Sex ratio remained close to parity across breeding sites and months, with female proportions generally ranging between 40% and 55% (**Fig. 7, Table 3**), and no consistent spatial or temporal trend. The close correspondence between larval and emergence patterns indicates that juvenile habitat quality strongly determines adult production, consistent with previous studies showing that resource limitation and environmental stress shape cohort output in aquatic insects [70, 71]. However, adult survival revealed more complex dynamics. The relatively high survival observed in B2 in October, despite reduced larval development and pupation, suggests a decoupling between early and late life stages. Such patterns are consistent with carry-over effects, where stressful larval environments reduce cohort size but may produce a subset of more stress-tolerant or phenotypically robust adults [72]. In mosquitoes, larval nutritional and physicochemical conditions are known to influence adult longevity and stress tolerance through persistent metabolic and endocrine adjustments [73, 74]. Together, these results support the idea that variation in juvenile habitat quality strongly structures adult population dynamics in organisms with complex life cycles [75], while also highlighting how carry-over effects can modulate, and in some cases decouple, adult performance from larval outcomes in heterogeneous urban environments.

Across treatments and experimental sessions, 65-71% of females laid eggs, with no significant differences in oviposition preference among B1, B2, and B3 or over sampling time points (all *p* > 0.28; **Fig. 8, table 4**). When considering the proportion of eggs laid in each breeding site versus the control, all sites were similarly attractive [B1: 80.4% (CI: 75%–84%); B2: 74% (68%–79%); B3: 79% (74%–84%); all *p* < 0.001], and no significant pairwise differences in preference strength were detected among breeding sites or across sampling time points within breeding sites (all adjusted *p* > 0.05). Thus, oviposition preference was high and temporally stable. Strikingly, females did not discriminate among B1, B2, and B3 despite strong differences in larval performance, indicating a decoupling between oviposition choice and offspring success. This suggests that females rely on multiple chemical and microbial cues when selecting oviposition sites, rather than simple proxies of habitat quality. Consistent with this, colorimetric and volatile profiles associated with microbial by⍰products and physicochemical signatures have been shown to influence *Aedes* spp. oviposition choices under semi⍰natural conditions [76, 77], potentially attracting females without necessarily indicating optimal fitness conditions. To determine whether early-life environmental differences extend beyond oviposition preference and larval performance, we evaluated the impact of breeding-site composition on egg fertility. Hatching rates were high and stable across breeding sites and months (mean = 82.4%; range: 33%-100%; **Fig. 9**), with no detectable effect on hatching success (all *p* > 0.34; site × month LRT *p* = 0.94). Hatching success remained consistent across conditions, typically ranging from 77% to 85%, suggesting that embryonic development is largely buffered against local environmental variation, likely due to maternal provisioning and the protective role of the eggshell [78]. Fertility was measured one month after oviposition by rehydrating desiccated viable eggs in breeding-site water collected at sampling time point T+2. In contrast, the effects of habitat quality became apparent only after hatching, during the larval stage, when development is highly dependent on microbial-mediated nutrition and oxygen dynamics [58, 79], rendering larvae more sensitive to physicochemical variation.

### Differential effects of abiotic and biotic parameters on mosquito fitness across developmental stages in breeding sites

To identify the environmental properties most closely associated with mosquito performance, we analyzed correlations between mosquito traits (life-history traits and sex ratio) and (i) abiotic characteristics of breeding sites and (ii) dominant microbial families present in these habitats over sampling time points. Only statistically significant correlations are presented (**Fig. 4–5**). Larval development time was associated exclusively with abiotic parameters, showing a positive correlation with paracetamol, whereas no significant association with microbial families was detected (**Fig. 4**). Because shorter development time generally reflects higher larval performance, this result suggests that paracetamol exposure is associated with delayed larval growth. Such delays may result from sublethal physiological or behavioral stress rather than direct acute toxicity. Recent work on *Ae. aegypti* has demonstrated that environmentally realistic concentrations of paracetamol can alter larval activity, learning, and memory, highlighting that micropollutants may disrupt mosquito performance even at low doses [80]. In contrast, larval survival was associated with both abiotic and biotic parameters. Among abiotic variables, survival was positively correlated with SO_4_^2−^, Na^+^, and Br^−^, but negatively correlated with NH_4_^+^ (**Fig. 4**). Pupation rate was influenced by the same compounds, consistent with the close relationship between larval viability and successful pupation. Some ions may indicate nutrient-rich conditions favourable to microbial productivity and larval nutrition, whereas elevated NH_4_^+^ concentrations are commonly associated with organic matter degradation, eutrophic conditions, and oxygen-depleted environments resulting from intense microbial decomposition [81, 82]. The negative correlation between NH_4_^+^ and larval survival is consistent with the known toxicity of ammonia in aquatic organisms, including mosquitoes, where elevated concentrations can impair performance and survival [82]. However, mosquito larvae are still frequently able to colonize ammonia-rich environments, reflecting their remarkable ability to tolerate and detoxify excess nitrogen. In particular, *Aedes* larvae regulate ammonia by excreting it and incorporating it into amino acids (glutamine, alanine and proline) and into compounds such as urea and uric acid [83–85], which helps them persist in organically enriched urban environments. Larval survival was also negatively correlated with the fungal family Cordycipitaceae and positively correlated with the bacterial family SC-I-84, indicating that microbial composition may further modulate habitat suitability (**Fig. 5**). Interestingly, Cordycipitaceae includes several entomopathogenic fungi such as *Beauveria bassiana*, which has repeatedly been shown to reduce larval survival, impair development, and alter oviposition behavior *in Ae. albopictus* [86, 87]. Adult emergence was exclusively associated with abiotic parameters, being positively correlated with Br^−^ and negatively correlated with N_2_O. While the ecological role of Br^−^ in mosquito breeding sites remains unclear, the negative association with N_2_O is consistent with previous studies linking elevated N_2_O concentrations to oxygen-depleted environments and intensified denitrification processes [88]. Duval et al. (2024) similarly reported an absence of larvae in urban containers with high N_2_O levels, suggesting that altered oxygen dynamics may reduce habitat suitability. Such conditions may be particularly detrimental during metamorphosis, as mosquito larvae partly rely on dissolved oxygen for respiration, and successful molting is impaired under hypoxic conditions [90]. Sex ratio was associated exclusively with abiotic parameters, being negatively correlated with Ca^2+^, CO_2_, and Mg^2+^ and positively correlated with paracetamol. These patterns likely reflect sex-specific sensitivity to environmental stress during larval development rather than direct effects on sex determination, as environmental stress is known to bias sex ratios through differential larval mortality [91–93]. Finally, adult survival was associated with both abiotic and biotic parameters, being positively correlated with Ca^2+^, adenine, CO_2_, and PeM15_fa, and negatively correlated with several bacterial families including Sphingomonadaceae, Microtrichaceae, and Trichocomaceae. These results suggest that adult longevity reflects carry-over effects from larval environmental conditions, where selective or stressful habitats may favour the emergence of more physiologically robust individuals [28, 94]. More broadly, the correlations observed with several bacterial families support the idea that microbial community composition can indirectly influence long-term mosquito fitness through effects on larval nutrition and development, as previously described [58, 73]. Overall, our results show that mosquito fitness in urban breeding sites is shaped by multiple environmental filters acting at different developmental stages. Abiotic conditions appear to primarily constrain larval development and metamorphosis, while microbial community composition is more strongly associated with larval survival and carry-over effects on adult performance.

In conclusion, our study demonstrates that urban mosquito breeding sites are highly heterogeneous environments where anthropogenic inputs structure physicochemical conditions and microbial communities across space and time. This environmental heterogeneity drives strong divergence in mosquito life-history traits while oviposition preference remains stable despite large differences in offspring performance, revealing a decoupling between habitat attractiveness and suitability. Overall, mosquito fitness is shaped by interacting abiotic and biotic filters that act differentially across developmental stages. These findings highlight the need to integrate environmental chemistry and microbial ecology to understand how urbanization shapes microbially mediated host– environment interactions and mosquito population dynamics.

## Acknowledgments

The authors acknowledge the Symbiotron platform from the FR BioEEnviS, as well as Angelo Jacquet for his help with the use of the insectary. They also thank the AME platform (University of Lyon, France) for access to gas and ionic chromatography facilities, as well as Danis Abrouk from the iBio platform (University of Lyon, France) for his help with sequence submission to GenBank. We also thank Imane El Drissi for her help in oviposition experiments.

## Author contributions

CVM and PL designed the study. CVM, PL, EM, LV, and AG collected samples and data, conducted the fitness and oviposition experiments, and analyzed bacterial composition. JG and AAMC performed the ionic and gaseous chromatography analyses. LW and AF conducted LC-HRMS suspect screening and LC-MS/MS quantification of micropollutants. AG, IT, and RC performed the statistical analyses. AG, CVM, and PL wrote the first draft of the manuscript. All authors reviewed and approved the final manuscript.

## Material and data availability

Supporting data for all results presented in this paper are contained within the manuscript and supplementary materials. The dataset analyzed during the current study and all R scripts used for statistical analyses and figure generation in this study are publicly available in the Zenodo repository, https://doi.org/10.5281/zenodo.20424249. DNA sequences of metabarcoding analyses are available in GenBank under accession numbers PRJEB113688 for 16S and PRJEB113689 for ITS.

---

**Figure S1.**
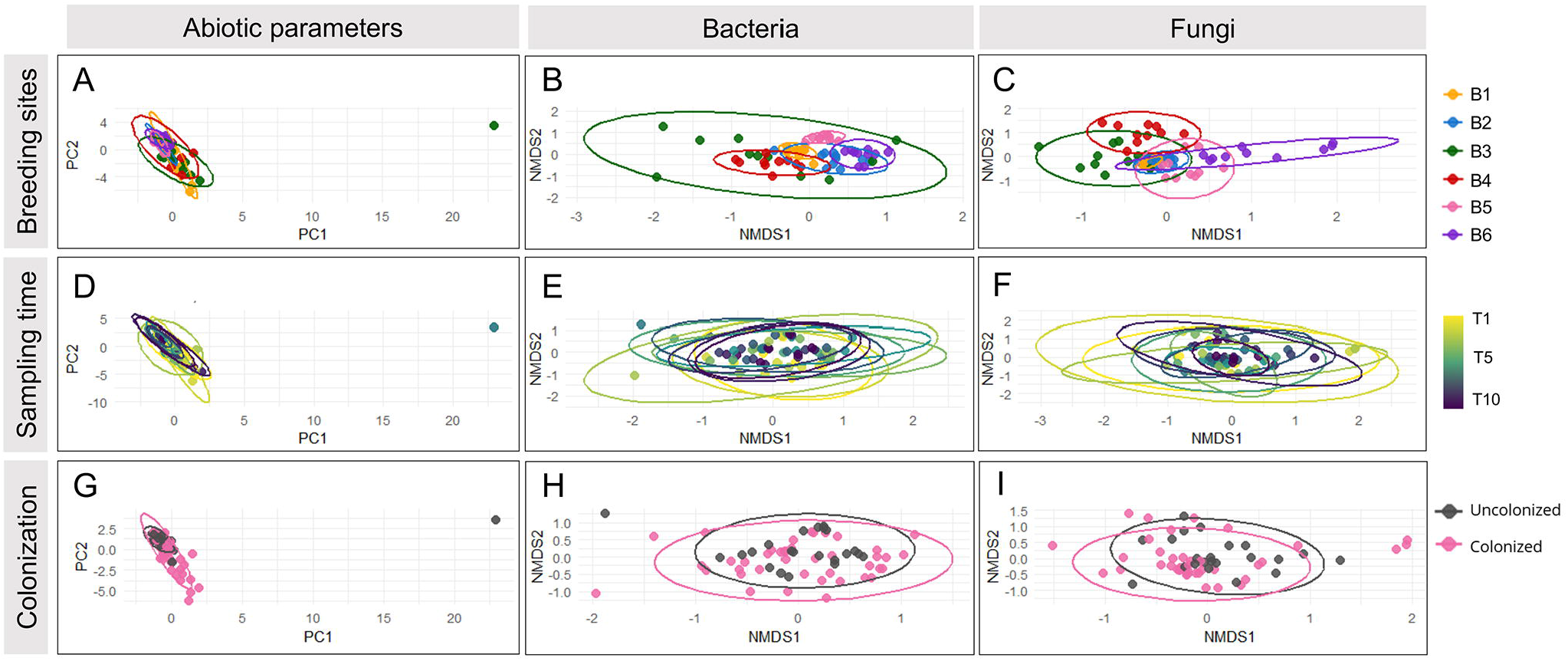
Non-metric multidimensional scaling (NMDS) and factor analysis of mixed data (FAMD) ordinations of physicochemical, bacterial, and fungal profiles across breeding-site identity, sampling time points, and colonization status. (A) FAMD ordination based on physicochemical parameters (ions, gases, micropollutants) colored by breeding-site identity (B1-B6). (B-C) NMDS ordinations based on Bray-Curtis dissimilarities of bacterial (16S) and fungal (ITS) community structures, respectively, colored by breeding-site identity (B1-B6). (D) FAMD ordination based on physicochemical parameters (ions, gases, micropollutants) colored by sampling time points (T1-T10). (E-F) NMDS ordinations based on Bray-Curtis dissimilarities of bacterial (16S) and fungal (ITS) community structures, respectively, colored by sampling time point (T1-T10). (G) FAMD ordination based on physicochemical parameters (ions, gases, micropollutants) colored by breeding-site colonization status (uncolonized vs colonized). (H-I) NMDS ordinations based on Bray-Curtis dissimilarities of bacterial (16S) and fungal (ITS) community structures, respectively, colored by breeding-site colonization status (uncolonized vs colonized).

**Figure S2.**
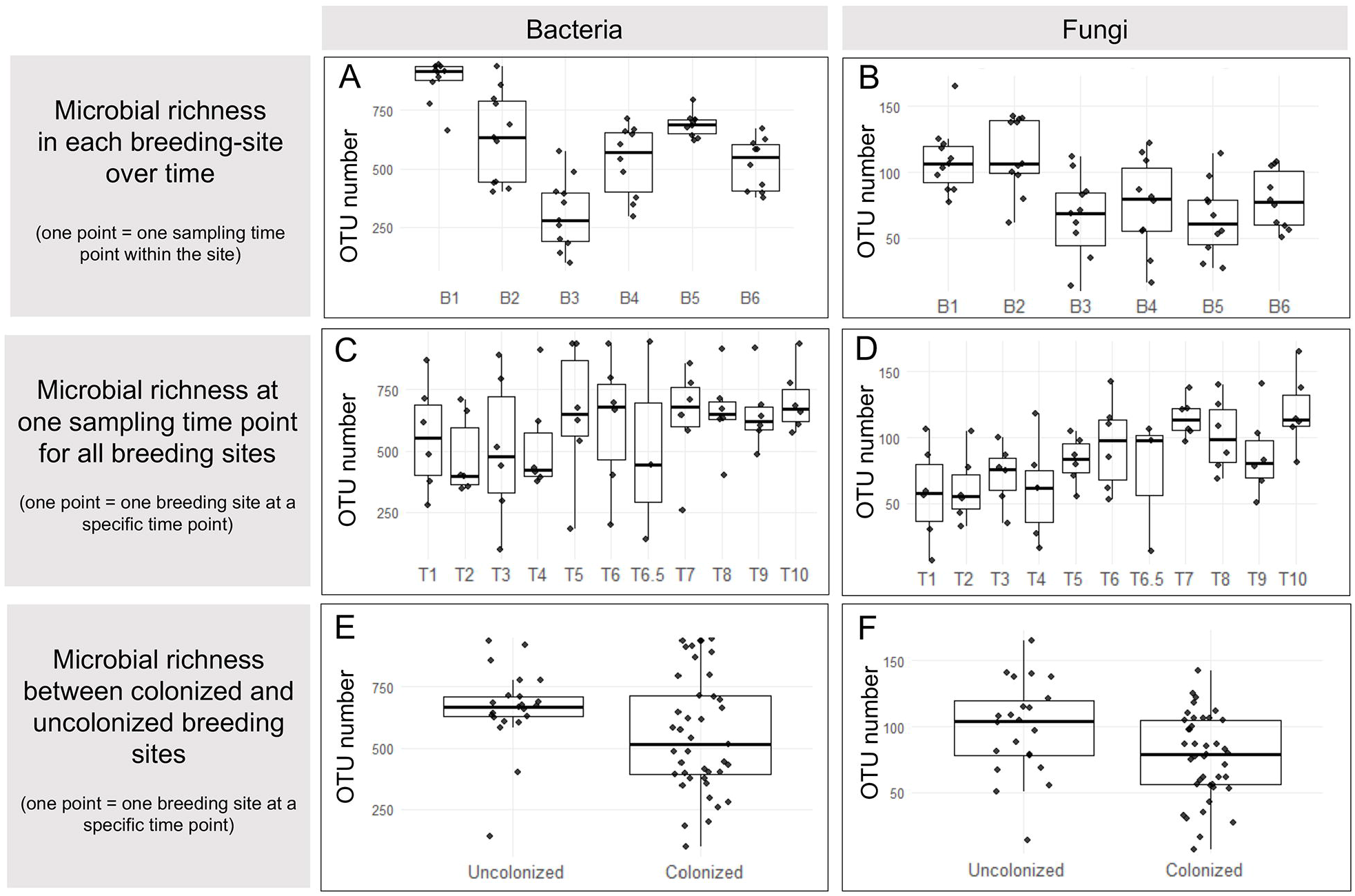
Bacterial and fungal richness across breeding site identity, sampling time points, and colonization status. (A-B) Boxplots representing the richness (total number of OTUs) of bacteria (A) and fungi (B) for each urban breeding site (B1-B6) over time. Each point corresponds to one sampling time point within a given breeding site. (C-D) Boxplots representing mean bacterial (C) and fungal (D) richness across all breeding sites by sampling time point (T1-T10). (E-F) Boxplots representing mean bacterial (E) and fungal (F) richness across all breeding sites and sampling time points according to colonization status (uncolonized/colonized). All boxplots display the median (thick line), interquartile range (bounds of the box), and whiskers (variability outside the interquartile range).

**Figure S3.**
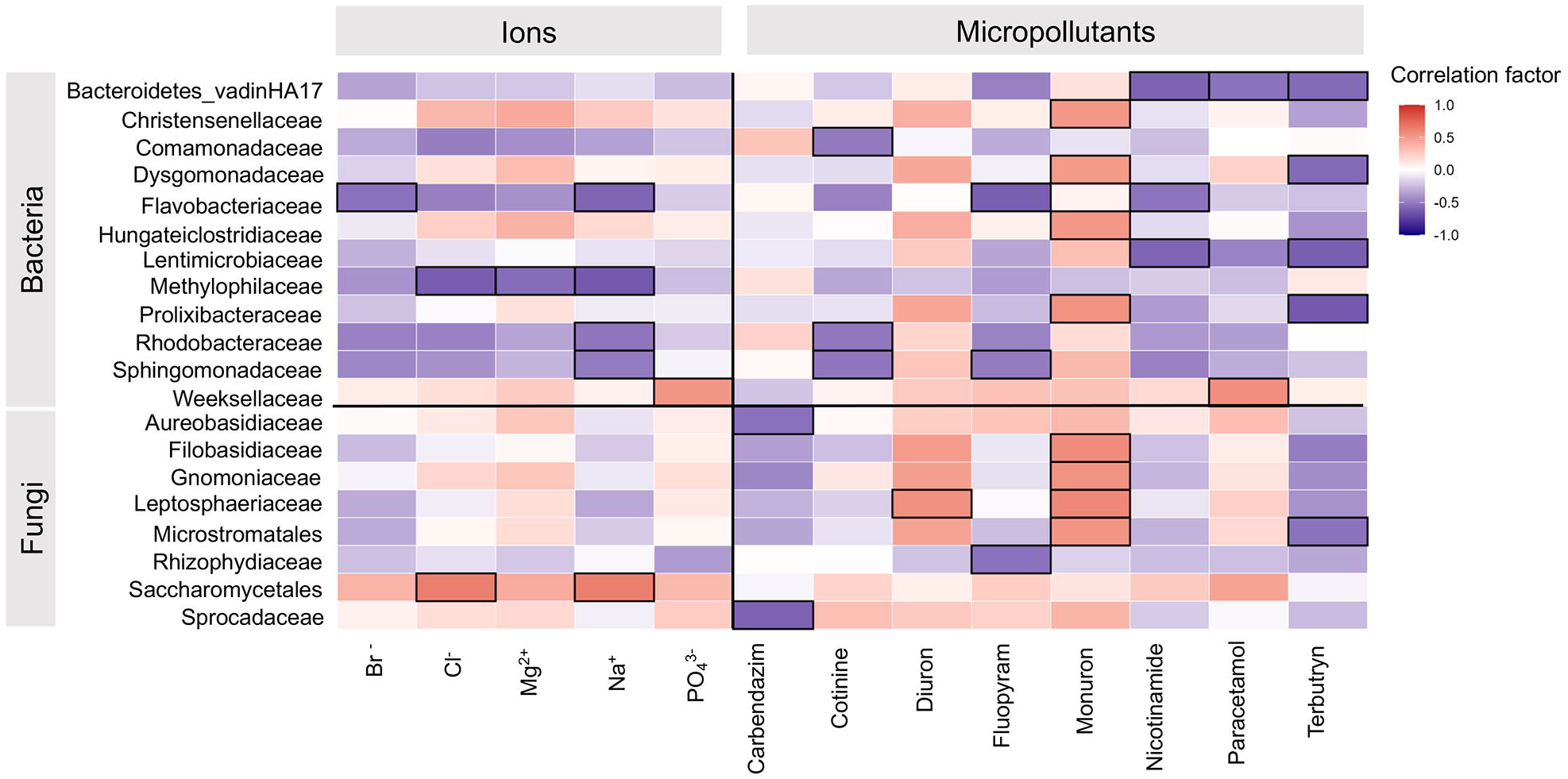
Significant correlations detected between dominant microbial families and selected abiotic factors. The most abundant bacterial and fungal families identified across breeding sites (B1-B6) were correlated with abiotic factors (pH, temperature, micropollutants, dissolved gases, and ions). Only the most statistically significant correlations are shown here; among the tested variables, only ions and micropollutants showed sufficiently strong associations with microbial families. Correlations with coefficients > 0.5 or < - 0.5 are outlined with black contour lines, with positive correlations colored in red and negative correlations in blue.

**Figure S4.**
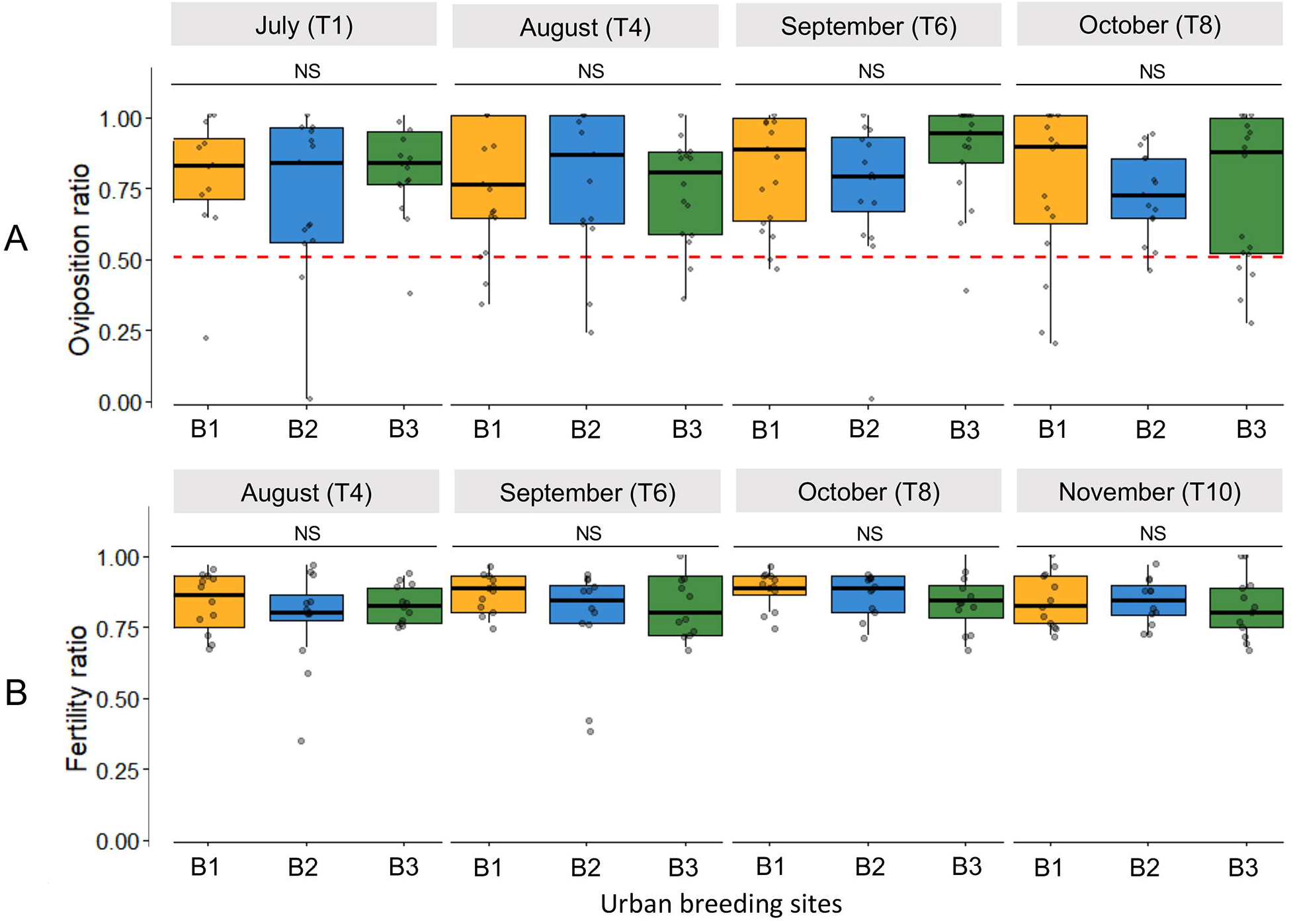
Temporal variation in the effects of urban breeding sites (B1-B3) on oviposition behavior and fertility. (A) Boxplots representing the ratio of eggs laid in the urban breeding site compared to the control beaker, across tested breeding sites (B1, B2, B3) and experimental time points (T1: July, T4: August, T6: September, T8: October). In this experiment, each gravid female (n=24) was individually placed into a net cage and allowed to choose between an artificial breeding site containing urban water and a control with sterile water. Non-responding females were included in the statistical analysis, whereas dead individuals were excluded. Oviposition behavior was analyzed using a two-step hurdle approach. Oviposition activation (whether females laid ≥1 egg) was assessed using a binomial model including breeding sites (B1-B3) and sampling time points as fixed effects. Oviposition preference (among females that laid eggs) was analyzed using a beta-binomial model comparing egg proportions between breeding sites and control waters. Preference was assessed by testing whether egg-laying proportions differed from 0.5 (no preference), with pairwise comparisons among breeding sites. For clarity, only differences between breeding sites within each sampling time point are shown. Temporal differences within each breeding site were also analyzed and are reported in Table S10. Statistical significance is shown above the boxplots with “*” indicating p < 0.05, “**” indicating p < 0.01, “***” indicating p < 0.001, and “NS” indicating *p >* 0,05. (B) Boxplots representing egg hatching success across breeding sites (B1, B2, B3) and experimental time points (T4: August, T6: September, T8: October, T10: November). Egg fertility was analyzed using beta regression with breeding sites, sampling time points, and their interaction as fixed effects, and plate identity as a random effect when appropriate. Proportions were transformed prior to analysis to avoid 0 and 1 values. Pairwise comparisons were adjusted using Tukey correction. As in panel (A), only differences between breeding sites within each sampling time point are displayed, while temporal differences within breeding sites are reported in Table S10. Statistical significance is indicated as described above. All boxplots display the median (thick line), interquartile range (bounds of the box), whiskers (variability outside the interquartile range), and outliers (points).

## References

1. O’Malley MA. The nineteenth century roots of ‘everything is everywhere’. Nat Rev Microbiol 2007;5:647–651. 10.1038/nrmicro1711

2. Rousk J, Bengtson P. Microbial regulation of global biogeochemical cycles. Front Microbiol 2014;5. 10.3389/fmicb.2014.00103

3. Cockell CS. Are microorganisms everywhere they can be? Environmental Microbiology 2021;23:6355–6363. 10.1111/1462-2920.15825

4. Mattoo R, B M S. Microbial roles in the terrestrial and aquatic nitrogen cycle—implications in climate change. FEMS Microbiol Lett 2023;370:fnad061. 10.1093/femsle/fnad061

5. Raza T et al. Unraveling the potential of microbes in decomposition of organic matter and release of carbon in the ecosystem. Journal of Environmental Management 2023;344:118529. 10.1016/j.jenvman.2023.118529

6. Cabanillas-Bojórquez LA et al. Microbial Production of Bioactive Compounds. In: Sarkar A, Ahmed IA (eds), Microbial products for future industrialization. Singapore: Springer Nature, 2023, 181–198.

7. Sadeghi J et al. Microbial community and abiotic effects on aquatic bacterial communities in north temperate lakes. Science of The Total Environment 2021;781:146771. 10.1016/j.scitotenv.2021.146771

8. Niu X et al. The combination of multiple environmental stressors strongly alters microbial community assembly in aquatic ecosystems. Journal of Environmental Management 2024;350:119594. 10.1016/j.jenvman.2023.119594

9. Bernardin JR et al. Environmental Stress Shapes Bacterial Community Structure and Function Through Interactive Abiotic Effects. Molecular Ecology 2025;34:e70102. 10.1111/mec.70102

10. David GM et al. Small freshwater ecosystems with dissimilar microbial communities exhibit similar temporal patterns. Molecular Ecology 2021;30:2162–2177. 10.1111/mec.15864

11. Vettorazzo S et al. From small water bodies to lakes: Exploring the diversity of freshwater bacteria in an Alpine Biosphere Reserve. Science of The Total Environment 2024;954:176495. 10.1016/j.scitotenv.2024.176495

12. Chaudhary A et al. Taxon-Driven Functional Shifts Associated with Storm Flow in an Urban Stream Microbial Community. mSphere 2018;3:10.1128/msphere.00194-18. https://doi.org/10.1128/msphere.00194-18

13. LaMartina EL, Mohaimani AA, Newton RJ. Urban wastewater bacterial communities assemble into seasonal steady states. Microbiome 2021;9:116. 10.1186/s40168-021-01038-5

14. Taipale SJ et al. Lowered nutritional quality of prey decrease the growth and biomolecule content of rainbow trout fry. Comparative Biochemistry and Physiology Part B: Biochemistry and Molecular Biology 2022;262:110767. 10.1016/j.cbpb.2022.110767

15. Johnson EB et al. Terrestrial Carbon Additions to Zooplankton Prey Influence Juvenile Estuarine Fish Growth. Environments 2023;10. 10.3390/environments10030050

16. Brown JJ et al. Metacommunity theory for transmission of heritable symbionts within insect communities. Ecology and Evolution 2020;10:1703–1721. 10.1002/ece3.5754

17. Sundaray JK et al. Metagenomic profiling of fish-associated microbiota: ecological perspectives from freshwater to marine environment—a review. Arch Microbiol 2026;208:105. 10.1007/s00203-025-04646-z

18. Apprill A. Marine Animal Microbiomes: Toward Understanding Host–Microbiome Interactions in a Changing Ocean. Front Mar Sci 2017;4. 10.3389/fmars.2017.00222

19. Pramanic A et al. Endophytic microbiota of floating aquatic plants: recent developments and environmental prospects. World J Microbiol Biotechnol 2023;39:96. 10.1007/s11274-023-03543-1

20. Gould AL et al. Microbiome interactions shape host fitness. Proc Natl Acad Sci U S A 2018;115:E11951–E11960. 10.1073/pnas.1809349115

21. O’Brien AM, Laurich JR, Frederickson ME. Evolutionary consequences of microbiomes for hosts: impacts on host fitness, traits, and heritability. Evol 2024;78:237–252. 10.1093/evolut/qpad183

22. Härer A et al. Associations Between Gut Microbiota Diversity and a Host Fitness Proxy in a Naturalistic Experiment Using Threespine Stickleback Fish. Molecular Ecology 2024;33:e17571. 10.1111/mec.17571

23. Rainford JL et al. Phylogenetic Distribution of Extant Richness Suggests Metamorphosis Is a Key Innovation Driving Diversification in Insects. PLOS ONE 2014;9:e109085. 10.1371/journal.pone.0109085

24. Guo T. Analysis of Life Cycle and Development Process of Holometabolous Insects. Molecular Entomology 2023;14.

25. Beutel RG, Goczał J, Pohl H. Evolutionary Adaptations in Larvae of Holometabola. Annual Review of Entomology 2026;71:149–168. 10.1146/annurev-ento-121423-013358

26. Jannot JE, Bruneau E, Wissinger SA. Effects of larval energetic resources on life history and adult allocation patterns in a caddisfly (Trichoptera: Phryganeidae). Ecological Entomology 2007;32:376– 383. 10.1111/j.1365-2311.2007.00876.x

27. Jiménez-Cortés JG, Serrano-Meneses MA, Córdoba-Aguilar A. The effects of food shortage during larval development on adult body size, body mass, physiology and developmental time in a tropical damselfly. Journal of Insect Physiology 2012;58:318–326. 10.1016/j.jinsphys.2011.11.004

28. Stoks R et al. Adaptive and Maladaptive Consequences of Larval Stressors for Metamorphic and Postmetamorphic Traits and Fitness. In: Costantini D, Marasco V (eds), Development Strategies and Biodiversity: Darwinian Fitness and Evolution in the Anthropocene. Cham: Springer International Publishing, 2022, 217–265.

29. Musolff A et al. Temporal and spatial patterns of micropollutants in urban receiving waters. Environmental Pollution 2009;157:3069–3077. 10.1016/j.envpol.2009.05.037

30. Hery L et al. Natural Variation in Physicochemical Profiles and Bacterial Communities Associated with Aedes aegypti Breeding Sites and Larvae on Guadeloupe and French Guiana. Microb Ecol 2021;81:93–109. 10.1007/s00248-020-01544-3

31. Duval P et al. Pollution gradients shape microbial communities associated with Ae. albopictus larval habitats in urban community gardens. FEMS Microbiology Ecology 2024;100:fiae129. 10.1093/femsec/fiae129

32. Braem S, Van Dyck H. Larval and adult experience and ecotype affect oviposition behavior in a niche-expanding butterfly. Behav Ecol 2023;34:547–561. 10.1093/beheco/arad022

33. Fowler EK et al. Female oviposition decisions are influenced by the microbial environment. j evol Biol 2025;38:379–390. 10.1093/jeb/voaf004

34. Malassigné S et al. Flower- and Water-Dwelling Yeasts Influence Nectar-Seeking and Oviposition Behavior in the Asian Tiger Mosquito with Distinct Volatile Organic Compound Profiles. J Chem Ecol 2026;52:3. 10.1007/s10886-025-01685-0

35. Gonzalez PV, González Audino PA, Masuh HM. Oviposition Behavior in Aedes aegypti and Aedes albopictus (Diptera: Culicidae) in Response to the Presence of Heterospecific and Conspecific Larvae. J Med Entomol 2016;53:268–272. 10.1093/jme/tjv189

36. Girard M et al. Microorganisms Associated with Mosquito Oviposition Sites: Implications for Habitat Selection and Insect Life Histories. Microorganisms 2021;9. 10.3390/microorganisms9081589

37. Mosquera KD et al. Multi-Omic Analysis of Symbiotic Bacteria Associated With Aedes aegypti Breeding Sites. Front Microbiol 2021;12. 10.3389/fmicb.2021.703711

38. Eastep NE, Albert RE, Anderson JR. Modulation of La Crosse Virus Infection in Aedes albopictus Mosquitoes Following Larval Exposure to Coffee Extracts. Front Physiol 2012;3. 10.3389/fphys.2012.00066

39. Antonelli P et al. Caught in a bad romance: Microbiota increases glyphosate toxicity in the Asian tiger mosquito Aedes albopictus. Environmental Pollution 2025;381:126651. 10.1016/j.envpol.2025.126651

40. Malassigné S et al. Environmental yeasts differentially impact the development and oviposition behavior of the Asian tiger mosquito Aedes albopictus. Microbiome 2025;13:99. 10.1186/s40168-025-02099-6

41. Raquin V et al. Variation in diet concentration and bacterial inoculum size in larval habitats shapes the performance of the Asian tiger mosquito, Aedes albopictus. Microbiome 2025;13:130. 10.1186/s40168-025-02067-0

42. Ihrmark K et al. New primers to amplify the fungal ITS2 region – evaluation by 454-sequencing of artificial and natural communities. FEMS Microbiol Ecol 2012;82:666–677. 10.1111/j.1574-6941.2012.01437.x

43. Lapworth DJ et al. Emerging organic contaminants in groundwater: A review of sources, fate and occurrence. Environmental Pollution 2012;163:287–303. 10.1016/j.envpol.2011.12.034

44. Herath JMMK et al. Breeding Habitat Preference of the Dengue Vector Mosquitoes Aedes aegypti and Aedes albopictus from Urban, Semiurban, and Rural Areas in Kurunegala District, Sri Lanka. J Trop Med 2024;2024:4123543. 10.1155/2024/4123543

45. Multini LC et al. The Influence of the pH and Salinity of Water in Breeding Sites on the Occurrence and Community Composition of Immature Mosquitoes in the Green Belt of the City of São Paulo, Brazil. Insects 2021;12. 10.3390/insects12090797

46. Scolari F et al. Exploring Changes in the Microbiota of Aedes albopictus: Comparison Among Breeding Site Water, Larvae, and Adults. Front Microbiol 2021;12. 10.3389/fmicb.2021.624170

47. Zhao SY et al. Microbiota Composition Associates With Mosquito Productivity Outcomes in Belowground Larval Habitats. Mol Ecol 2025;34:e17614. 10.1111/mec.17614

48. Allison SD, Martiny JBH. Colloquium paper: resistance, resilience, and redundancy in microbial communities. Proc Natl Acad Sci U S A 2008;105 Suppl 1:11512–11519. 10.1073/pnas.0801925105

49. Duval P et al. Pollution gradients shape microbial communities associated with Ae. albopictus larval habitats in urban community gardens. FEMS Microbiol Ecol 2024;100:fiae129. 10.1093/femsec/fiae129

50. Fang K et al. Environmental stressors drive fungal community homogenization and diversity loss in plateau freshwater lakes. BMC Microbiol 2025;25:438. 10.1186/s12866-025-04144-8

51. Duguma D et al. Bacterial Communities Associated with Culex Mosquito Larvae and Two Emergent Aquatic Plants of Bioremediation Importance. PLOS ONE 2013;8:e72522. 10.1371/journal.pone.0072522

52. Rogy P, Srivastava DS. Terrestrial subsidies and light affect an aquatic micro-ecosystem in unexpected ways. Freshwater Biology 2024;69:879–893. 10.1111/fwb.14252

53. Burke C et al. Bacterial community assembly based on functional genes rather than species. Proc Natl Acad Sci U S A 2011;108:14288–14293. 10.1073/pnas.1101591108

54. Shade A et al. Fundamentals of microbial community resistance and resilience. Front Microbiol 2012;3:417. 10.3389/fmicb.2012.00417

55. Backhaus T, Faust M. Predictive environmental risk assessment of chemical mixtures: a conceptual framework. Environ Sci Technol 2012;46:2564–2573. 10.1021/es2034125

56. Di Nica V et al. Toxicity of Quaternary Ammonium Compounds (QACs) as single compounds and mixtures to aquatic non-target microorganisms: Experimental data and predictive models. Ecotoxicology and Environmental Safety 2017;142:567–577. 10.1016/j.ecoenv.2017.04.028

57. Smith TP et al. High-throughput characterization of bacterial responses to complex mixtures of chemical pollutants. Nat Microbiol 2024;9:938–948. 10.1038/s41564-024-01626-9

58. Coon KL et al. Mosquitoes rely on their gut microbiota for development. Molecular Ecology 2014;23:2727–2739. 10.1111/mec.12771

59. Luis P et al. Aedes albopictus mosquitoes host a locally structured mycobiota with evidence of reduced fungal diversity in invasive populations. Fungal Ecology 2019;39:257–266. 10.1016/j.funeco.2019.02.004

60. Staley ZR, Harwood VJ, Rohr JR. A synthesis of the effects of pesticides on microbial persistence in aquatic ecosystems. Crit Rev Toxicol 2015;45:813–836. 10.3109/10408444.2015.1065471

61. Souza RS et al. Microorganism-Based Larval Diets Affect Mosquito Development, Size and Nutritional Reserves in the Yellow Fever Mosquito Aedes aegypti (Diptera: Culicidae). Frontiers in Physiology 2019;10.

62. Ponnusamy L et al. Oviposition responses of Aedes mosquitoes to bacterial isolates from attractive bamboo infusions. Parasites Vectors 2015;8:486. 10.1186/s13071-015-1068-y

63. Pesce S, Margoum C, Montuelle B. In situ relationships between spatio-temporal variations in diuron concentrations and phototrophic biofilm tolerance in a contaminated river. Water Research 2010;44:1941–1949. 10.1016/j.watres.2009.11.053

64. Rosi EJ et al. Urban stream microbial communities show resistance to pharmaceutical exposure. Ecosphere 2018;9:e02041. 10.1002/ecs2.2041

65. Fleeger JW, Carman KR, Nisbet RM. Indirect effects of contaminants in aquatic ecosystems. Sci Total Environ 2003;317:207–233. 10.1016/S0048-9697(03)00141-4

66. Relyea R. The impact of insecticides and herbicides on the biodiversity and productivity of aquatic communities. Ecol Appl. Ecological Applications 2005;15:618–627. 10.1890/1051-0761(2006)016%5B2027:TIOIAH%5D2.0.CO;2

67. Riaz MA et al. Impact of glyphosate and benzo[a]pyrene on the tolerance of mosquito larvae to chemical insecticides. Role of detoxification genes in response to xenobiotics. Aquat Toxicol 2009;93:61–69. 10.1016/j.aquatox.2009.03.005

68. Perumal V et al. A review of entomopathogenic fungi as a potential tool for mosquito vector control: A cost-effective and environmentally friendly approach. Entomological Research 2024;54:e12717. 10.1111/1748-5967.12717

69. Muturi EJ et al. Association between fertilizer-mediated changes in microbial communities and Aedes albopictus growth and survival. Acta Trop 2016;164:54–63. 10.1016/j.actatropica.2016.08.018

70. Yan J, Kibech R, Stone CM. Differential effects of larval and adult nutrition on female survival, fecundity, and size of the yellow fever mosquito, Aedes aegypti. Front Zool 2021;18:10. 10.1186/s12983-021-00395-z

71. Mackay AJ et al. Larval diet and temperature alter mosquito immunity and development: using body size and developmental traits to track carry-over effects on longevity. Parasit Vectors 2023;16:434. 10.1186/s13071-023-06037-z

72. Crean AJ, Monro K, Marshall DJ. Fitness consequences of larval traits persist across the metamorphic boundary. Evolution 2011;65:3079–3089. 10.1111/j.1558-5646.2011.01372.x

73. Dickson LB et al. Carryover effects of larval exposure to different environmental bacteria drive adult trait variation in a mosquito vector. Sci Adv 2017;3:e1700585. 10.1126/sciadv.1700585

74. Kang DS et al. Larval stress alters dengue virus susceptibility in Aedes aegypti (L.) adult females. Acta Trop 2017;174:97–101. 10.1016/j.actatropica.2017.06.018

75. Gerber R et al. Body stores of emergent aquatic insects are associated with body size, sex, swarming behaviour and dispersal strategies. Freshwater Biology 2022;67:2161–2175. 10.1111/fwb.14003

76. Allgood DW, Yee DA. Oviposition preference and offspring performance in container breeding mosquitoes: evaluating the effects of organic compounds and laboratory colonisation. Ecol Entomol 2017;42:506–516. 10.1111/een.12412

77. Montini P, Fischer S. Oviposition site selection and subsequent offspring performance of Aedes aegypti in short- and long-term detritus accumulation conditions. Acta Tropica 2024;255:107222. 10.1016/j.actatropica.2024.107222

78. Mosquera KD et al. Egg-laying by female Aedes aegypti shapes the bacterial communities of breeding sites. BMC Biol 2023;21:97. 10.1186/s12915-023-01605-2

79. Silberbush A, Abramsky Z, Tsurim I. Dissolved oxygen levels affect the survival and developmental period of the mosquito Culex pipiens. Journal of Vector Ecology 2015;40:425–427. 10.1111/jvec.12186

80. Dessart M, Lazzari CR, Guerrieri FJ. Acute and chronic sublethal chemical pollution affects activity, learning and memory in mosquito larvae. J Exp Biol 2026;229:jeb250493. 10.1242/jeb.250493

81. Ward BB. Nitrification and ammonification in aquatic systems. Life Support Biosph Sci 1996;3:25– 29.

82. Walker ED. Toxicity of Sulfide and Ammonium to Aedes triseriatus Larvae (Diptera: Culicidae) in Water-Filled Tree Holes and Tires. J Med Entomol 2016;53:577–583. 10.1093/jme/tjw032

83. von Dungern P null, Briegel H. Protein catabolism in mosquitoes: ureotely and uricotely in larval and imaginal Aedes aegypti. J Insect Physiol 2001;47:131–141. 10.1016/s0022-1910(00)00096-2

84. Scaraffia PY et al. Differential ammonia metabolism in Aedes aegypti fat body and midgut tissues. J Insect Physiol 2010;56:1040–1049. 10.1016/j.jinsphys.2010.02.016

85. Durant AC, Donini A. Development of Aedes aegypti (Diptera: Culicidae) mosquito larvae in high ammonia sewage in septic tanks causes alterations in ammonia excretion, ammonia transporter expression, and osmoregulation. Sci Rep 2019;9:19028. 10.1038/s41598-019-54413-6

86. Kirsch JM, Tay J-W. Larval Mortality and Ovipositional Preference in Aedes albopictus (Diptera: Culicidae) Induced by the Entomopathogenic Fungus Beauveria bassiana (Hypocreales: Cordycipitaceae). Journal of Medical Entomology 2022;59:1687–1693. 10.1093/jme/tjac084

87. Tawidian P, Kang Q, Michel K. The Potential of a New Beauveria bassiana Isolate for Mosquito Larval Control. J Med Entomol 2023;60:131–147. 10.1093/jme/tjac179

88. Naqvi SW et al. Increased marine production of N2O due to intensifying anoxia on the Indian continental shelf. Nature 2000;408:346–349. 10.1038/35042551

89. Duval P et al. Pollution gradients shape microbial communities associated with Ae. albopictus larval habitats in urban community gardens. FEMS Microbiology Ecology 2024;100:fiae129. 10.1093/femsec/fiae129

90. Alvarez-Costa A et al. Challenging Popular Belief, Mosquito Larvae Breathe Underwater. Insects 2024;15. 10.3390/insects15020099

91. Farjana T, Tuno N, Higa Y. Effects of temperature and diet on development and interspecies competition in Aedes aegypti and Aedes albopictus. Medical and Veterinary Entomology 2012;26:210–217. 10.1111/j.1365-2915.2011.00971.x

92. Cardôso HCB et al. Effects of predation by the copepod Mesocyclops ogunnus on the sex ratios of mosquito Aedes albopictus. Hydrobiologia 2013;705:55–61. 10.1007/s10750-012-1379-3

93. Bara JJ, Montgomery A, Muturi EJ. Sublethal effects of atrazine and glyphosate on life history traits of Aedes aegypti and Aedes albopictus (Diptera: Culicidae). Parasitol Res 2014;113:2879–2886. 10.1007/s00436-014-3949-y

94. Monaghan P. Early growth conditions, phenotypic development and environmental change. Philos Trans R Soc Lond B Biol Sci 2008;363:1635–1645. 10.1098/rstb.2007.0011

