## Supplementary material for "From urban runoff to mosquito success: spatiotemporal microbial assembly in larval water habitats under anthropogenic stressors"

**Supplementary Materials:** **additional information concerning the Materials and Methods section of the paper “From urban runoff to mosquito success: spatiotemporal microbial assembly in larval water habitats under anthropogenic stressors**”

**Study sites and sampling design**

At each sampling time point, water was subdivided into three subsamples: 250 mL for ionic and gaseous chromatography collected in a DURAN^®^ graduated laboratory bottle (Schott), 400 mL for micropollutants analyses collected in a high-density polyethylene bottle (VWR), and 50 mL for microbial community analyses collected in a Falcon tube (Greiner Bio-One). Samples for micropollutant and metabarcoding analyses were stored at -20°C, whereas samples for ionic and gaseous analyses were stored at +4°C until processing.

**Physicochemical characterization of breeding site waters**

For dissolved gases, thawed aliquots were equilibrated in airtight flasks with a 50 mL helium headspace at 60°C for 3 h under shaking (150 rpm). Gas concentrations were measured using a micro gas chromatograph (μGC R990, SRA Instruments) equipped with two columns connected to a thermal conductivity detector (TCD): an MS5A stainless-steel column (10 m × 0.25 mm, 30 μm BF) for O₂ and N₂, and a PPQ column (10 m × 0.25 mm, 8 μm) for CH₄, N₂O, and CO₂. Calibration gas mixtures ranged from 0-14,000 ppm (CO₂), 0-50,000 ppm (N₂O), 0-100,000 ppm (CH₄), 0-79% (N₂), and 0-21% (O₂). Inorganic ions were quantified by ion chromatography (AQUION, Thermo Fisher Scientific) coupled to an autosampler (AS-AP 120) with conductivity detection. Anions were separated at 30°C on an AG9-HC guard column and an AS9-HC analytical column (4 × 250 mm) using 9 mM Na₂CO₃ at 1 mL min^-1^ and a 4 mm carbonate suppressor (AERS 500). Cations were analyzed at 40°C on a CG-19 guard column and a CS-19 analytical column (4 × 250 mm) using 30 mM methanesulfonic acid at 1 mL min^-1^ with a 4 mm suppressor (CDRS 600). Target ions included F⁻, Cl⁻, NO₂⁻, Br⁻, NO₃⁻, PO₄³⁻, SO₄²⁻, Na⁺, NH₄⁺, K⁺, Mg²⁺, and Ca²⁺. Calibration curves (0-500 mg L^-1^) were prepared from 1 g L^-1^ stock solutions in ultrapure water.

Organic micropollutants were analyzed using a suspect screening workflow coupled with vacuum assisted evaporative concentration and UHPLC–HRMS. Briefly, 100 mL of water samples were filtered through 0.7 μm glass fiber filters (Millipore IT30 142 HW system), and spiked with 20 μL of a 2 mg L^-1^ mixture of 6 isotopically labelled internal standards. Water samples were concentrated at 55 °C using a vacuum-assisted evaporation system (Syncore® Analyst, BÜCHI Labortechnik AG, Switzerland). Parallel evaporation from up to 6 glass vials down to a residual volume of approximately 1 mL lasted 120 min. Concentrated extract were transferred to vials and centrifuged at 10,000 rpm for 10 min before being split into two fractions. Suspect screening analyses were performed using an Ultimate 3000 UHPLC (Thermo Fisher Scientific®) coupled to a QToF mass spectrometer (Maxis Plus, Bruker®) with an ESI source. Two chromatographic separation were applied : (i) reverse-phase separation on an Intensitysolo C18 column, Bruker® (2.2 μm, 100 × 2.1 mm) with a KrudKatcher Ultra in-line filter guard (Phenomenex®) at 30°C using a 5 μL injection volume and (ii) HILIC separation on a SeQuant ZicHilic column, Merck® (3.5 µm, 100 x 2.1 mm). The reverse-phase mobile phase consisted of (A) water:methanol 99:1 and (B) methanol, both containing 5 mM ammonium formate and 0.01% formic acid, whereas the HILIC gradient used (A) water:acetonitrile 50:50 and (B) water:acetonitrile 95:5, both containing 10 mM ammonium formate. HRMS acquisition was carried out in positive ionization mode (capillary 3600 V; end plate offset 500 V; nebulizer 3 bar N₂; drying gas 9 L min^-1^; drying temperature 200°C) with external exact-mass calibration using sodium formate/acetate clusters. Data were acquired in data-independent acquisition (DIA) mode with alternating MS and MS/MS scans over m/z 50–1000 at 40 eV collision energy. Data preprocessing and suspect screening were conducted in TASQ v1.4 (Bruker®) using the PesticideScreener 2.1 and ToxScreener 2.1 databases (≈1200 pesticides and ≈800 pharmaceuticals) and an in-house database (≈120 polar compounds). Identification criteria included ±5 ppm exact mass tolerance, ±0.3 min retention time deviation, mSigma < 20 for isotope pattern matching, and ±20 ppm fragment ion tolerance. More details are available in Fildier et al. 2025.

After the identification of micropollutants, a targeted quantification approach was performed by direct injection (10 µL) of the water samples, using an UHPLC H-Class (Waters®) coupled to a Xevo TQ-S (Waters®) triple quadrupole mass spectrometer. Multiple Reaction Monitoring (MRM) acquisition was conducted in positive ESI mode (capillary 3000 V; source temperature 150 °C; nebulizer 7 bar N_2_; drying gas N_2_ 900 L h^-1^ at 550 °C). The chromatographic separation was achieved on a Luna Omega Polar column C18 (3 µm 100 x 2.1 mm, Phenomenex®) using a gradient of (A) water with 0.01% formic acid and (B) methanol. .

**Environmental DNA extraction, metabarcoding, and sequence processing**

Samples were centrifuged at 16,000 × g for 10 min at 4°C and pellets were resuspended in 250 μL pre-heated CTAB lysis buffer (2% CTAB, 1.4 M NaCl, 100 mM Tris-HCl pH 8.0, 20 mM EDTA, and 0.2% β-mercaptoethanol). Samples were incubated at 60°C for 1 h with gentle mixing, followed by RNase A treatment (0.4 mg per reaction) for 5 min at 37°C. Nucleic acids were purified by sequential extraction using phenol:chloroform:isoamyl alcohol (25:24:1, v/v/v), followed by chloroform:isoamyl alcohol (24:1, v/v). eDNA was precipitated with isopropanol, washed with 75% ethanol, air-dried, and resuspended in RNase-free water.

Paired-end reads were merged with VSEARCH, and unmerged reads were discarded. Sequences outside the expected amplicon length ranges (250-300 bp for 16S and 200-500 bp for ITS) were removed. Filtered reads were clustered into Operational Taxonomic Units (OTUs) using SWARM with an aggregation distance of one nucleotide (d = 1), which iteratively aggregates highly similar amplicon variants. Chimeric sequences and low-abundance OTUs (< 0.005% of total reads) were subsequently removed. Taxonomic assignment was performed using marker-specific reference databases. Bacterial OTUs were classified against the SILVA database (v138.1), whereas fungal OTUs were assigned using the UNITE Fungi database (v8.3) with the RDP classifier. The UNITE eukaryotic reference (v8.2) was additionally used to improve fungal taxonomic resolution. To account for potential contamination introduced during eDNA extraction, sequence counts were corrected using negative extraction controls, as previously described (Antonelli et al. 2025).

**Oviposition assays**

A laboratory colony of *Ae. albopictus* (*AealbVB*), established from individuals collected in 2017 in Villeurbanne and Pierre-Bénite (France), was used for all experiments. Egg hatching was induced overnight at 28 °C under reduced pressure (-20 mm Hg). Larvae were reared at 28 °C in plastic trays containing dechlorinated water under a 16:8 h light:dark photoperiod and fed daily with a powdered diet consisting of 75% tropical fish food (TetraMin, Tetra) and 25% yeast tablets (Biover) until pupation. Newly emerged adults were transferred to BugDorm cages (32.5 × 32.5 × 32.5 cm) and maintained in environmental chambers (Panasonic MLR-352) set at 28 °C, 80% relative humidity, and a 16:8 h light:dark cycle, with continuous access to a 10% sucrose solution. Females aged 5-10 days were offered a blood meal using an artificial membrane feeding system (Hemotek Ltd) fitted with pig intestine membranes and filled with defibrinated sheep blood (Thermo Fisher Scientific). Fully engorged females were maintained in clean cages for 48 h to allow egg maturation.

For each oviposition assay, blood-fed gravid females were placed individually in BugDorm cages (20 × 20 × 20 cm) containing two 50 mL glass beakers: one with 40 mL filtered breeding-site water and the other with 40 mL sterile water as a control. Each beaker contained a strip of blotting paper moistened with sterile water to serve as an oviposition substrate. Each breeding-site water and sampling time point combination included 24 biological replicates. Beaker positions were randomized within cages to avoid positional bias. A small container with cotton soaked in sterile 10% sucrose solution was provided as a food source. Females were allowed to oviposit for four days under controlled environmental conditions (28 °C, 80% relative humidity, 16:8 h light:dark cycle).

To generate axenic larvae, viable eggs were gently detached from blotting papers using a toothbrush and transferred into sterile 50 mL conical tubes with filter caps (Greiner Bio-One). Eggs were rinsed with 50 mL of sterile water (Gibco) and surface-sterilized by successive immersions (5 min each) in 70% ethanol, 3% sodium hypochlorite, and 70% ethanol, followed by three rinses in sterile water. Egg hatching was induced overnight at 28°C under vacuum conditions (−20 mm Hg) in the presence of 50 μL sterile 10% liquid fish food. Ten axenic first-instar larvae (L1) were transferred into each well of sterile six-well plates (Starlab) containing a standardized food plug composed of 5% (w/v) crushed tropical fish flakes. For each breeding site (B1-B3), five plates were prepared, and five additional plates containing sterile water served as axenic controls. Plates were incubated at 28°C in complete darkness, and larval development was monitored daily for 21 days to record mortality and pupation time. Pupae were collected individually under sterile conditions and placed into 1.5 mL microtubes with slightly loosened caps to allow adult emergence. Survival during larval development and metamorphosis was recorded to estimate pupation rate, adult emergence success, and sex ratios. Newly emerged adults were sexed, pooled by plate, and transferred to BugDorm cages (20 x 20 x 20 cm) maintained at 28°C, 80% relative humidity, and a 16:8 h light:dark cycle, with continuous access to a 10% sucrose solution. Adult longevity was monitored every 2-3 days for up to 60 days, and deceased individuals were sexed post-mortem. Egg fertility was evaluated from eggs obtained during oviposition assays. After collection, eggs were dried for one month, counted under a binocular magnifier, and visually inspected. Eggs that appeared cracked, collapsed, or otherwise damaged were classified as non-viable and excluded. To assess hatching success, viable eggs were rehydrated approximately one month after oviposition, reflecting the natural temporal scale of the mosquito life cycle. Eggs were placed in water collected from the corresponding breeding site at sampling time point T+2 rather than in the original oviposition water. Two six-well plates were prepared per experimental condition. Eggs laid in breeding-site water were rehydrated in water from the same site, whereas eggs laid in sterile water were rehydrated in sterile water (control). Twenty-four viable eggs were gently transferred into each well using a sterile cotton swab for each breeding-site condition. Egg deposition in control conditions was occasionally low, resulting in reduced sample sizes (9–10 eggs per well depending on the sampling time point). Each well received 6 mL of breeding-site or sterile water, and plates were incubated in darkness at 28°C for 72 h before hatched larvae were counted.

**Statistical analyses**

Abiotic variables were standardized and explored using principal component analysis (PCA) to assess environmental variation among breeding sites and sampling time points. Microbial community composition was analyzed using non-metric multidimensional scaling (NMDS) based on Bray–Curtis dissimilarities calculated from relative abundance OTU tables derived from bacterial (16S) and fungal (ITS) datasets. Alpha diversity (richness and diversity indices) was analyzed using generalized linear models (GLMs) with breeding site, sampling time point, and colonization status as predictors, and significance was assessed via likelihood ratio tests. Community structure and abiotic variables were further evaluated using one-factor PERMANOVA analyses testing the effects of breeding site, sampling time point, and colonization status independently. Oviposition behavior was analyzed using a two-step hurdle approach: oviposition activation (whether females laid ≥1 egg) and oviposition preference (for females that laid eggs) were assessed using a binomial model including breeding site (B1-B3) and sampling time point as fixed effects and a beta-binomial model comparing egg proportions between breeding-site and control water, respectively. Preference was assessed by testing whether egg-laying proportions differed from 0.5 (no preference), with Tukey-adjusted pairwise comparisons among breeding sites. Larval development time was analyzed using accelerated failure time (AFT) models with right-censoring, comparing Weibull, log-normal, and log-logistic distributions using Akaike’s Information Criterion (AIC). The best-supported model included breeding site, sampling time point, and their interaction. Pairwise contrasts were estimated using marginal means with Tukey-adjusted *P*-values. Larval survival, pupation probability, and adult emergence were analyzed using binomial generalized linear mixed models with breeding site, sampling time point, and their interaction as fixed effects, and, where appropriate, well, plate, or cage identity as random effects to account for non-independence among observations. Mosquito sex ratio was analyzed using binomial models of female and male counts per cage, with breeding site, sampling time point, and their interaction as fixed effects, and final inference was based on a binomial GLM after the cage random effect was found to be negligible, with Tukey-adjusted pairwise comparisons. Adult survival was analyzed using AFT models with model selection based on AIC and confirmed using a Cox proportional hazards model accounting for cage clustering. Egg fertility (hatching success) was analyzed using beta regression with breeding site, sampling time point, and their interaction as fixed effects, and plate identity was included as a random effect when appropriate, with proportions transformed to avoid 0 and 1 values prior to modeling, while pairwise comparisons were adjusted using Tukey or Holm corrections depending on the model structure and comparison type. To assess associations between abiotic and biotic parameters and mosquito traits, we used Spearman’s rank correlation coefficient. This non-parametric approach was chosen because it captures monotonic relationships without assuming linearity or normality of the data. Statistical significance was assessed using a permutation test with 1,000 permutations, and correlations were considered statistically significant at *p* < 0.05. To improve robustness and reduce the influence of sparse or poorly characterized taxa, several preprocessing steps were applied. First, OTUs were aggregated at the family level. Second, families with insufficient information were removed by retaining only those with at least 5 non-zero values among the 12 observations. For biotic variables, we additionally required a minimum of 1,000 total reads per family to avoid disproportionate influence of rare taxa with low read counts. Finally, unclassified families were removed from the analyses.

**Bibliography**

Antonelli, Pierre, Laurent Vallon, Edwige Martin, et al. 2025. « Caught in a bad romance: Microbiota increases glyphosate toxicity in the Asian tiger mosquito *Aedes albopictus* ». *Environmental Pollution* 381 (septembre): 126651. https://doi.org/10.1016/j.envpol.2025.126651.

Fildier A, Zerbini G, Wiest L, Gentil A, Valiente Moro C, Vulliet E. Suspect screening based on reversed-phase and HILIC LC-HRMS of micropollutants in tiger mosquito urban breeding sites. JFSM 2025, Jun 2025, Montpellier, France. <https://hal.science/hal-05039866v1>
